# Objective assessment of visual attention in toddlerhood

**DOI:** 10.1101/2023.04.04.534573

**Authors:** E. Braithwaite, V. Kyriakopoulou, L. Mason, A. Davidson, N. Tusor, N. Harper, M. Earl, S. Datoo-Partridge, A. Young, A. Chew, S. Falconer, Joseph V Hajnal, M.H. Johnson, C. Nosarti, A.D. Edwards, E.J.H. Jones

**Author notes:** Joint first authors (E. Braithwaite, V. Kyriakopoulou, L. Mason). Joint last authors (A.D. Edwards, E.J.H. Jones).

## Abstract

Visual attention is an important mechanism through which children learn about their environment, and individual differences could substantially shape later development. Eyetracking provides a sensitive and scalable tool for assessing visual attention that has potential for objective assessment of child development, but to date the majority of studies are small and replication attempts are rare. This study investigates the feasibility of a comprehensive eye-tracking assessment of visual attention and introduces a shared data resource for the scientific community. Data from eight eyetracking tasks were collected from 350 term-born (166 females) 18-month-olds recruited as neonates http://www.developingconnectome.org/). Analyses showed expected condition effects for seven of eight tasks (*p*-values from <.001 to .04), an important indication of replicability. Consistent with some theoretical models of visual attention, structural equation modelling indicated participants’ performance could be explained by two factors representing social and non-social attention. Comprehensive eye-tracking batteries can objectively measure individual differences in core components of visual attention in large-scale toddlerhood studies. This is the first large-scale comprehensive study to present high-quality normative eye-tracking data from a large task battery in toddlers and make them freely available to the scientific community.

## Introduction

The first three years of life are critical for brain development (Fox et al., 2010) and have a significant impact on later quality of life and economic contributions (Knudsen et al., 2006). Most current approaches to large-scale measurement of early brain and cognitive development either use indirect measures (e.g. head circumference), or behavioural assessments. This severely limits our capacity to rapidly and objectively track early cognitive development as behavioural measures require skilled assessors, rely on children’s verbal comprehension and motor skills, and may not be translatable to different contexts and cultures. To provide more direct assessment of cognitive development, one highly promising method is eye-tracking. Eye-tracking is a non-invasive technology that can provide exquisite temporal and spatial resolution on a child’s direction of gaze, and can be largely automated to produce scalable measures of individual differences in visual attention in early development. Humans are a highly visual species, and control information input through eye movements and visual attention is a particularly crucial modality for learning in early development. Eye-tracking tasks can be conducted without the need for complex verbal instruction and do not rely upon children’s comprehension ability or motor skills (Karatekin, 2007; Richmond & Nelson, 2009). Eye-tracking limits potential researcher bias as it minimises the role of the researcher and provides a more direct and objective measure of children’s processing (Karatekin, 2007; Sasson & Elison, 2012). Additionally, compliance rates are usually high since there are no requirements for the child to wear any equipment, nor the need to interact with anyone unfamiliar to them, enabling a broader community of children to successful participate (Karatekin, 2007). This is particularly important in toddlerhood, where the challenges involved in working with children with limited verbal and executive functioning skills but strong motor skills have limited progress in studying neurocognitive development in the second and third years of life.

Many developmental studies have utilised eye-tracking to investigate attention, but these have typically focussed on comparing group differences in performance on a single eye-tracking task or a battery of tasks in small size cohorts (B. M. Hood & Atkinson, 1993; Hutchinson & Turk-Browne, 2012; van Baar et al., 2020). Single task studies have yielded important findings about how bias for faces changes over development (Leppänen, 2016), how orienting to stimuli differs under conditions of competition (B. M. Hood & Atkinson, 1993) and how memory guides attention (Hutchinson & Turk-Browne, 2012). However, much less is known about whether eyetracking can provide appropriate measures of individual differences, and whether tasks can be combined effectively into larger batteries.

Recently, a multiple-task battery showed reasonable reliability and validity for use with toddlers, though sample size was limited (n = 12 in the reliability sample) (van Baar et al., 2020). To use eye-tracking as a large-scale, comprehensive assessment of development, acquisition and robustness of task performance when multiple tasks are combined must be examined.

Using a comprehensive battery of visual attention tasks also allows investigators to move beyond metrics extracted from single tasks and to test whether profiles of attention across the battery fit are consistent with theoretical models of visual attention. In designing eye-tracking tests of attention, many investigators have distinguished between endogenous and exogenous control of attention (Connor et al., 2004; Sarter et al., 2001). Endogenous control involves executive attention systems that select goal-relevant actions by resolving conflict between competing inputs or impulses (Amso & Scerif, 2015; Miller & Cohen, 2001). Conversely, exogenous control is stimulus-driven, and relies primarily on ‘bottom-up’ mechanisms.

Although most work on the development of attention has employed non-social stimuli, there has been increasing recent interest in the construct of ‘social attention’, the motivation to attend to social stimuli such as people and faces. This is of particular interest given its perturbation in the emergence of neurodevelopmental conditions such as autism (Klin et al., 2015). A rich historical literature indicates that infants orient to faces from birth (Farroni et al., 2005) and show preference for face-like stimuli (versus non-face configurations), direct gaze (versus averted gaze or eyes-closed gaze) and biological motion (versus inverted or scrambled) throughout development (Shultz et al., 2018). However, the intersection between behaviours labelled as ‘social attention’ and the exogenous and endogenous attention systems described above remains unclear (see Braithwaite et al., 2020 for review). Possibly, attention to social stimuli represents a specific case of attention to object features or ’what’ forms of attention. Alternatively, social stimuli may gain attention due to the interplay between social motivation and other attentional systems (Chevallier et al., 2012; Dawson et al., 2005). Whilst such substructures have been examined longitudinally and through examination of condition differences, it remains unclear whether there are meaningful individual differences in separable subdomains of visual attention.

### The current study

In this paper we present data collected from a large cohort of 18-month-old children tested on a large battery of eye-tracking tasks designed to measure both exogenous and endogenous attentional control in toddlerhood. Tasks were selected for inclusion based on previous literature, and relevance to assessing exogenous, endogenous and social attention (Elsabbagh et al., 2009, 2013), Kidd et al. (2012) Wass et al. (2011) Kaldy et al. (2011); (Saez de Urabain et al., 2017); (Võ et al., 2012). Several tasks are also present in other large-scale data collection efforts in younger infants (Falck-Ytter et al., 2021; Hessels & Hooge, 2019; Jones et al., 2019, 2019), providing important opportunities for synergies. We examine the robustness of these tasks to combined delivery and examine whether derived metrics can provide independent estimates of core domains of visual attention. For each task, we examined data quality, attrition rates, and the presence or absence of expected condition effects. We then tested the fit of a theoretical model of visual attention to the pattern of data across tasks using structural equation modelling, to determine whether a smaller set of underlying constructs explained individual differences in visual attention within this cohort. All data and extracted metrics presented in this study are available to the scientific community as part of the developing Human Connectome Project scheduled data release (http://www.developingconnectome.org/).

## Materials and Methods

### 2.1 Ethics

The study was approved by the West London & GTAC Research ethics committee (REC: 14/LO/1169). Written informed consent was obtained from the parents or legal guardians of all participating children. The data are available from the National Institute of Mental Health Data Archive (https://nda.nih.gov/), under Developing Human Connectome Project (dHCP) Collection 3955 (https://nda.nih.gov/edit_collection.html?id=3955). All access requests are reviewed and handled by the NDA.

### 2.2. Cohort

The cohort included 350 term-born children (184 males, 166 females) recruited from low-risk pregnancies as part of the Developing Human Connectome Project (DHCP, http://www.developingconnectome.org/). Children aged 18 months who had been recruited to the study as neonates were invited to attend an eye-tracking assessment and a neurodevelopmental assessment at the Centre for the Developing brain in St Thomas’ hospital. Exclusion criteria were preterm birth (<37 weeks) and the presence of a sibling with a diagnosis of ASD. Cohort demographics are presented in Table S1.

**Table 1:** Mean, standard deviation and range for demographic information

| Cohort Demographics |  | Mean | (SD) | Range |
| --- | --- | --- | --- | --- |
| Gestational age at birth (weeks) |  | 39.9 | (±1.3) | (37-43) |
| Age at assessment (months) |  | 18.3 | (±1.1) | (17-24) |
| Sex |  |  |  |  |
|  | Males (%) | 184 | (53%) |  |
|  | Females (%) | 166 | (47%) |  |
| Bayley Scores |  |  |  |  |
|  | Cognitive composite score | 100 | (±10.9) | (55-125) |
|  | Language composite score | 97.3 | (±15.3) | (50-153) |
|  | Motor composite score | 102 | (±9.3) | (67-130) |
| IMD quintile groups (%) |  |  |  |  |
|  | 1 | 7.7 % |  |  |
|  | 2 | 9.8 % |  |  |
|  | 3 | 18.1 % |  |  |
|  | 4 | 34.7 % |  |  |
|  | 5 | 29.8 % |  |  |
| Ethnicity |  |  |  |  |
|  | Maternal |  |  |  |
|  | White | 59.2 % |  |  |
|  | Black | 16.1 % |  |  |
|  | Asian | 8.9 % |  |  |
|  | Chinese | 1.4 % |  |  |
|  | Other | 14.4 % |  |  |
|  | Paternal |  |  |  |
|  | White | 62.4 % |  |  |
|  | Black | 17.2 % |  |  |
|  | Asian | 8.9 % |  |  |
|  | Chinese | 0.0 % |  |  |
|  | Other | 11.5 % |  |  |
| English first language |  |  |  |  |
|  | Both parents | 51.3 % |  |  |
|  | Neither parent | 30.3 % |  |  |
|  | One parent | 17.5 % |  |  |

### 2.3 Eye tracking data acquisition

Eye tracking data were acquired from a Tobii TX-300 (Tobii AB, Sweden) at a sampling rate of 120Hz, with children sitting approximately 60cm from the 23” screen (58.42cm x 28.6cm, 52.0° x 26.8° @ 60cm, native resolution of 1920 x 1080 pixels and an aspect ratio of 16:9). Stimuli were presented on Apple (Apple Inc., USA) Macbook Pro computers, using our custom-written stimulus presentation framework (Task Engine, sites.google.com/site/taskenginedoc/), running in Matlab using Psychtoolbox 3 (Brainard, 1997; Kleiner et al., 2007) and the GStreamer library (gstreamer.freedesktop.org) for video decoding (SM1.1 for more details).

### 2.4. Tasks

At the start of the eye tracking assessment the experimenter positioned each participant in front of the eye tracker. Online feedback was given to allow a position to be chosen as close as possible to the centre of the eye tracker head box, to maximise data quality. An automatic five-point calibration was then performed. At the beginning of each trial, a gaze-contingent fixation stimulus was presented in the centre of the screen; when gaze fell upon this stimulus the trial started. All tasks except for the Gap-Overlap began when the participant fixated a gaze-contingent central fixation stimulus, at a size of 3cm x 3cm (2.86° x 2.86° at 60cm viewing distance). Tasks were administered in blocks that were intermixed with each other and distributed throughout the battery. SM1.2 provides additional task details.

#### 2.4.1 Gap-Overlap

##### 2.4.1.1. Stimulus presentation

Trials were presented in blocks of 12. Each trial started with the onset of a central stimulus (CS), a cartoon image of an analogue clock accompanied by an alerting sound. When the infant fixated the CS, after a 600-700ms wait period the peripheral stimulus (PS) was presented at 21.7° of visual angle on the left or right of the screen (random). In the baseline condition the CS disappeared concurrent with the appearance of the PS. In the overlap condition the CS continued to be presented for the duration of the rest of the trial. In the gap condition the CS was removed from the screen 200ms before the PS onset. The PS was a cartoon cloud that appeared on either the left or the right side of the screen and was accompanied by a sound, 3cm (2.86°) from the edge, rotating at 500° per second until fixated by the participant. A reward stimulus was then presented at the location of the PS for 1000ms.

##### 2.4.1.2. Data extraction

Mean SRTs (time of first sample to enter the PS – time of PS onset) were calculated for gap, overlap and baseline conditions (<u>GO-Gap-SRT</u>; <u>GO-Baseline-SRT</u>; <u>GO-Overlap-SRT</u>) separately, using only valid trials (see SM1.2.1.2). For examination of condition differences, disengagement scores (<u>GO-Disengagement)</u> were computed as Overlap-SRT minus Baseline-SRT and facilitation scores (<u>GO-Facilitation)</u> were computed as Baseline-SRT minus Gap-SRT. Reaction times are expected to be fastest in the gap then baseline then overlap condition (Elsabbagh et al., 2009).

#### 2.4.2 Non-social contingency

##### 2.4.2.1. Stimulus presentation

This task consisted of three blocks of 19 trials. Blocks were distributed across the battery in a randomised order but constrained such that the 60% condition was always presented within the first two blocks. At the start and end of each block, a static picture of four balls was presented for a fixed period of 5 seconds. Between these was a set of trials. In each trial, participants were first presented with a fixation stimulus (hummingbird, 3cm x 3cm, 2.86° x 2.86°) that remained on screen until it was fixated by the infant. The fixation stimulus was replaced by a display of four balls (4.5cm, 2.3°), one in each corner of the screen, 3cm (2.9°) from each edge. Participants then saccaded to one of the four balls (within an AOI 50% bigger than the ball, subtending 4.3° degrees of visual angle); the ball they selected became their ‘chosen’ ball for that trial. The ‘reward’ for choosing a ball was that one of the balls would become animated; in the 100% reward block it was always the ball they chose that was animated; in the 0% reward block it was never the ball they chose (one of the other balls was randomly selected) and in the 60% reward block on 60% of the trials it was the ball they chose that was animated (and on the remaining 40%, a randomly chosen ball was animated). Rewards lasted 1000ms.

##### • 2.4.2.1. Data extraction

Offline: Saccadic reaction time to select a ball was computed as the difference between the frame on which the four balls appeared and the first sample to enter one of the four AOIs and averaged within a block. Fixation times to return to the central stimulus at the beginning of each trial were an index of engagement and computed as the difference between the frame on which the fixation stimulus appeared and the first sample to enter the fixation stimulus and averaged within a block. Trials were excluded if the reward was not played (to ensure the child had selected a ball) or if reaction time was less than 150ms (to avoid trials where children were looking at a ball when the trial started); skipped trials were also excluded.

Key dependent variables were saccadic reaction time to select a ball with removing reaction times > 2000ms (“zone outs”; <u>NSC-100-SRT-NoZone</u>, <u>NSC-60-SRT-NoZone; NSC-0-SRT-NoZone</u>); and fixation times to return to the central stimulus at the beginning of each trial (<u>NSC-100-FixRT</u>, <u>NSC-60-FixRT; NSC-0-FixRT</u>). Reaction times are expected to be fastest in the 60% condition because it is moderately predictable (Kidd et al., 2012).

#### 2.4.3 Reversal learning (‘Cognitive Control’)

##### 2.4.3.1. Stimulus presentation

This task was based on Wass et al. (2011). Two purple 17cm by 13cm (16.1° x 12.5° @ 60cm) rectangles were presented on the left and right of the screen (1.5cm, or 1.43° from the outermost edge). These remained on screen until either 1) either one of the rectangles was fixated by the participant, or 2) 2000ms had elapsed. At this point, one of the rectangles was replaced by video of the same dimensions, showing a 2s clip of the animated children’s TV programme Thomas the Tank Engine. After the 2s clip had played, the screen became blank and the next trial began with another gaze-contingent fixation stimulus.

In the first (non-scored) trial, the side that the child chose to look at first was recorded; if no side was chosen after 2000ms it was determined randomly. On the following set of 8 trials (termed the “learning phase”), the video was always presented on the opposite side to that chosen on the first trial. This learning phase ended after either a) the child made three anticipatory saccades to the correct side of the screen, or b) eight trials (excluding the first, non-scored, trial) had been presented. The task then entered the “reversal phase” where the correct side was reversed and an additional nine trials were presented. The first trial of the reversal phase was not scored, but instead served to indicate (implicitly) to the child that the correct side had been reversed.

##### 2.4.3.2. Data extraction

AOIs were placed around the location of each of the rectangles (within one of which the video played) and dilated by 2° to account for poor calibration. Accuracy was calculated as *number of correct trials/ total number of trials where an anticipation occurred* and saccadic reaction times were logged, then averaged across valid trials. Trials were considered invalid if SRT was less than 300ms or if no antisaccade was made. Participants were excluded from analysis if they made fewer than 2 antisaccades (regardless of correctness) with valid SRTs per phase (learning/reversal), per block.

Key dependent variables were pre-switch accuracy (<u>CC-Pre-Acc)</u> and reaction time <u>(CC-Pre-SRT);</u> we also assessed task performance using post-switch accuracy (<u>CC-Post-Acc)</u>, and reaction time (and <u>CC-Post-SRT</u>) but because these were not completed by all participants and were dependent on performance in the pre-switch phase (missing not at random) they were not included in later modelling. Accuracy is expected to be lower and reaction times are expected to be slower during the pre-switch phase (in line with Wass et al., 2011).

#### 2.4.4 Working memory

##### 2.4.4.1. Stimulus presentation

Images of two theatre stages with a lowered curtain were drawn on either side of the screen (16.0cm x 23.4cm, 15.2° x 22.1° @ 60cm). The stages remained presented through the trial. Over 500ms a toy appeared and dropped from the top to the vertical centre of the screen. Once the participant fixated the toy, the curtains on both the left and right stages lifted over 400ms. Over the next 750ms the toy moved to one of the (randomly chosen) stages. The toy remained motionless for 200ms before spinning for 400ms to engage attention at its stopping point on the stage. Over 400ms both stage curtains lowered, hiding the toy.

A central fixation was then presented until it was fixated by the participant; at this point it paused for 200ms before spinning for 200ms to engage attention and then disappeared. The task then waited for the participant to fixate one of the two curtains (defined as gaze being over the curtain for minimum 100ms); the choice was coded to be either correct or incorrect, depending upon whether or not the chosen curtain was hiding the toy. Over the next 400ms the chosen curtain was raised, revealing either the toy or an empty stage, depending upon whether the participant chose the correct side. If correct, the toy spun for 400ms as a reward before dropping off the bottom of the screen. The curtain then lowered. The reaction time (relative to the offset of the spinning central fixation stimulus) to choose a curtain, and the correctness of the choice, were recorded.

##### 2.4.4.2. Data extraction

Accuracy was calculated as number of correct trials/ total number of valid trials. Saccadic reaction times were computed as the difference between the timestamps of the presentation of the first two rectangles and the first sample to enter a rectangle AOI, averaged across valid trials. Trials were considered invalid and were discarded if 2000ms elapsed without either curtain being fixated. The key dependent variables were accuracy of selecting the correct curtain (<u>WM-Acc)</u> and saccadic reaction times overall (<u>WM-SRT-All);</u> we additionally compared reaction times for each of correct and incorrect responses (<u>WM-SRT-Acc</u> and <u>WM-SRT-Inacc</u>). Infants were expected to achieve above chance-level accuracy (Daehler et al., 1976; Hofstadter & Reznick, 1996).

#### 2.4.5 Visual search

##### 2.4.5.1. Stimulus presentation

Search displays consisted of three different items; red apple (target, 4.57cm x 4.57cm, 4.3° x 4.3°), blue apple (colour distractor, 4.57cm x 4.57cm, 4.3° x 4.3°) and a red slice of an apple (an elongated rectangle, cropped from the full apple image, shape distractor, 1.12cm x 6.67cm, 1.1° x 6.4°). These stimuli were used to produce two trail types, feature search and conjunction search trials.

Stimuli in feature search trials varied on only one dimension, either colour or shape (e.g. a red apple surrounded by blue apples, or a red full apple surrounded by red slices). The set size in these trials was always 9, with one target stimulus and 8 distractors. Conjunction search trials consisted of an equal number of both colour and shape distractors, and the set size was either 9 or 13. In total there were six possible configurations of trial, two feature search trials and four conjunction search trials.

Stimuli were arranged on screen algorithmically. All stimuli were required to be within a circular region located at the centre of screen with a diameter of 27.2cm (25.5° @ 60cm). The target stimulus was then positioned at a random point within this circle, ensuring that it was not within 6cm (5.7° @ 60cm) of the centre of the screen, where it would overlap with the central fixation stimulus presented at the start of each trial. Next, each distractor stimulus was placed at a random location within the circle, ensuring that no stimulus (target or distractor) overlapped the spatial location of any other stimulus.

Each trial began with an animated central stimulus unique to this task. The target stimulus (a red apple) “flew” into the screen from one edge (left, top, right, bottom – chosen randomly) over 800ms, ending at the centre of the screen. When the participant fixated the apple, it faded into the background colour of the screen over 750ms, and then the full array was drawn. Note that the location of the target amongst the distractors was not the central location of this attention-getter, indeed the target was never presented within 6cm of this location. The array was presented for 4000ms or until the participant fixated the target stimulus, after which the target span as a visual reward for 1300s until the trial ended.

##### 2.4.5.2. Data extraction

Offline: AOIs were drawn around the target and distractors. Accuracy was calculated as the number of trials on which the infant fixated the target (apple) before the animation was automatically triggered/ the number of trials administered. Saccadic reaction time to find the target on each trial was the difference between the time at which the search slide was presented and the time of the first gaze sample in the target AOI for each condition. Trials were excluded if reaction time was less than 150ms or if the trial had been skipped.

The key dependent variables are accuracy (<u>VS-S9-Acc</u>, <u>VS-C9-Acc</u>, <u>VS-C13-Acc)</u> and the saccadic reaction time to find the target on each trial (<u>VS-S9-SRT</u>, <u>VS-C9-SRT</u>, <u>VS-C13-SRT</u>). Reaction times are expected to be fastest in the single search, then conjunctive 9-item search, then the conjunctive 13-item search (in line with Gerhardstein & Rovee-Collier, 2002).

#### 2.4.6 Face Pop-out

##### 2.4.6.1. Stimulus presentation

Infants were presented with a series of six annular visual arrays each composed of five objects in different locations on the screen (Gliga, Elsabbagh, Andravizou & Johnson, 2009; Hendry et al, 2018). Each array contained: a face with direct gaze, a visual ‘noise’ image generated from the same face presented within the array by randomising the phase spectra of the face whilst keeping the amplitude and colour spectra constant to act as a control for the low-level visual properties of the face stimuli (Halit, Csibra, Volein & Johnson, 2004), a bird, a car and a mobile phone.

##### 2.4.6.2. Data extraction

Areas-of-interest (AOIs) masks were placed around each stimulus. For each AOI, the proportion of samples within it was calculated by *number of samples in AOI/ number of valid (non-missing) samples.* Contiguous runs of samples within an AOI were identified and the mean proportion looking time and peak look duration to each AOI were calculated across valid trials only (see SM1.2.5.2.).

Peak look to each AOI is the duration of the longest look to that AOI during each trial averaged across the number of valid trials. Key dependent variables are the proportion of trials on which the infants first look to the face divided by the number of trials with a valid initial look (reflective of social orienting, <u>Pop-Face-First</u>), percent looking to faces (<u>Pop-Face-Pct</u>, reflective of general interest (Elsabbagh et al., 2013)), and two key comparative AOIs for examination of the success of the manipulation, selected because they are the best visual control for faces (scrambled face or ‘Noise’) or are an object of interest (Car); (<u>Pop-Car-Pct</u>, <u>Pop-Noise-Pct</u>); peak look to each AOI (reflective of sustained attention(Gui et al., 2020) <u>Pop-Face-Peak</u>, <u>Pop-Car-Peak</u>, <u>Pop-Noise-Peak</u>). Percentage looking and looking times are expected to be greater to faces than comparison stimuli (Gliga et al., 2009).

#### 2.4.7 Dancing ladies

##### 2.4.7.1. Stimulus presentation

Each trial consisted of the presentation of one of six videos (20-25s duration each, 25 fps, 1920×1080 resolution) of three women dancing with objects. Videos were designed to be semi-naturalistic. Three videos were played in their native format and three matched videos were visually-scrambled such that the social information was degraded.

##### 2.4.7.2. Data extraction

Offline: AOIs were hand traced around the faces and objects on a frame-by-frame basis using Motion (Apple Inc, USA) software, and were approximately 14.5° x 33.5° @ 60cm. Samples were assigned to AOIs and interpolated across gaps of < 200ms preceded and succeeded by the same AOI. Percent attention to each AOI was computed as *number of samples in each AOI/ total number of valid samples* for that video and averaged across videos within each condition; peak look to each AOI was defined as the longest run of samples within one AOI during each video, averaged across videos within each condition. Data was excluded if there were fewer than 25% valid samples for each video. Key dependent variables were percent attention to faces, <u>Dance-Soc-Face-Pct</u>, <u>Dance-Scr-Face-Pct</u>) and peak look (<u>Dance-Soc-Face-Peak</u>, <u>Dance-Scr-Face-Peak</u>). <u>I</u>n addition, as a comparator we examined percent attention to the object (<u>Dance-Soc-Object-Pct, Dance-Scr-Object-Pct</u>); peak look to the object (<u>Dance-Soc-Object-Peak</u>, <u>Dance-Scr-Object-Peak</u>). Looking times are expected to be greater to faces than comparison stimuli (Frank et al., 2014; Gliga et al., 2009).

#### 2.4.8 50 Fifty faces

##### 2.4.8.1 Stimulus presentation

Originating from the “50 People, One Question” project (Krolak, 2011), infants watched a video comprised of street interviews in English with a number of people (41s, 1280px x 720px, 25fps): We removed the soundtrack of the original video and replaced it with classical music in order not to introduce linguistic confounds.

##### 2.4.8.2 Data extraction

Offline: AOIs were hand traced around the faces, bodies and background people on a frame-by-frame basis using Motion (Apple Inc, USA) software. Samples were assigned to AOIs and interpolated across gaps of <200ms preceded and succeeded by the same AOI. Percent attention to each AOI was computed *as number of samples in each AOI/total number of valid samples for the video*; peak look to each AOI was defined as the longest run of samples within one AOI during the video. Key dependent variables were percent attention to faces (<u>50Face-Face-Pct</u>); peak look to faces (<u>50Face-Face-Peak</u>). Data was excluded if there were fewer than 25% valid samples for the video. To examine whether social attention was greater than to other elements of the scene, we also examined looking to the background (<u>50Face-Background-Pct, 50Face-Background-Peak).</u> Looking times are expected to be greater to faces than comparison stimuli (Frank et al., 2014; Gliga et al., 2009).

### 2.5. Data quality assessment

In addition to task-specific data quality metrics (duration of valid data extracted from free viewing tasks and number of trials available from trial-based tasks), we computed two measures of the general quality of the eye-tracking data across the session. Accuracy (the spatial displacement of recorded gaze from the point fixated) and precision (variability in consecutive samples on the same fixation point) were extracted as proxies of general eye-tracking quality across the session (see SM1.3 for details).

### 2.6. Statistics

We first examined a range of variables pertinent to the valid acquisition of data from each task. These included the percentage of children who provided valid data; whether key metrics significantly varied with data quality (operationalised as having a correlation coefficient >= .2 or <= -.2); and whether the expected pattern of condition differences was elicited by the tasks administered in combination (SM1.4 for details). Expected condition differences were assessed using Analysis of Variance tests (ANOVAs) or one-tailed t-tests where appropriate and where we had a strong directional hypothesis. Analyses were repeated after children with too few valid trials were removed (SM2.3) and additional analyses were conducted to assess the impact of task-specific data quality metrics on condition effects (see SM2.2.2 & SM2.2.3 for details). Finally, we used a hypothesis-driven SEM modelling approach to examine the underlying structure of visual attention (using lavaan (Best, 2020)). Selected scores from individual tasks were z-scored (see Figure 3); we initially selected items to fit a theoretically motivated three-factor model (representing social attention, and exogenous and endogenous shifting) with equal numbers of items (three items from three tasks each). We used a factor analysis to decompose key data quality metrics (Accuracy, Precision, Gap valid trial numbers, reversal learning trial numbers, proportion lost samples in the popout, dancing ladies and fifty faces) into two factors representing accuracy and precision, and data quantity; these factors were then entered into the confirmatory factor analysis as covariates.

## Results

### 3.1 Feasibility of the battery

Overall, both data retention and data quality were good, indicating that the current eye-tracking battery was successfully implemented.

#### 3.1.1 Data retention

Retention of data was good (Table 2). A high proportion of children (at least 70%) reached and provided at least some valid data for each task in the battery. For all but the non-social contingency task, over 90% of those who reached the task provided enough data to be included in condition effect analyses and for all tasks fewer than 10% of these participants were excluded after cut-off criteria were applied. Tasks towards the end of the battery were typically reached by fewer participants, though a high percentage (over 90%) of the children who reached these tasks did provide usable data.

**Table 2:** Data retention number and rates for each task. All percentages are based on children who had enough data for inclusion on the highest yield variable

| Task | N and % of children who got to task in battery (includes any participant who provided any valid data at all) |  | N and % with enough data for whole-group condition effect analysis |  | N and % with enough data for cut-off analysis |  |
| --- | --- | --- | --- | --- | --- | --- |
| Gap-Overlap | 337 | 96.3% | 332 | 94.9% | 302 | 86.3% |
| Non-Social contingency | 333 | 95.1% | 253 | 72.3% | 253 | 72.3% |
| Reversal Learning | 323 | 92.3% | 323 | 92.3% | 312 | 89.1% |
| Working Memory | 332 | 94.9% | 331 | 94.6% | 311 | 88.9% |
| Visual Search | 343 | 98.0% | 341 | 97.4% | 318 | 90.9% |
| Face pop-out | 256 | 73.1% | 256 | 73.1% | 246 | 70.3% |
| Dancing Ladies | 251 | 71.7% | 249 | 71.1% | 226 | 64.6% |
| Fifty faces | 246 | 70.3% | 246 | 70.3% | 235 | 67.1% |

#### 3.1.2 Data quality

Data quality was generally good (Fig. 1). Accuracy and precision were 1.7 (*SD* = 0.8) and 1.5 (*SD* = 0.5) degrees respectively, with AOIs typically ranging from 4.3-39 degrees, indicating that the quality of the eye-tracking data was generally sufficient for the task design. Despite cleaning and validation procedures, accuracy and precision did associate significantly with individual differences in key variables across many tasks (SM2.2.1, tables S4 & S5).

**Figure 1:**
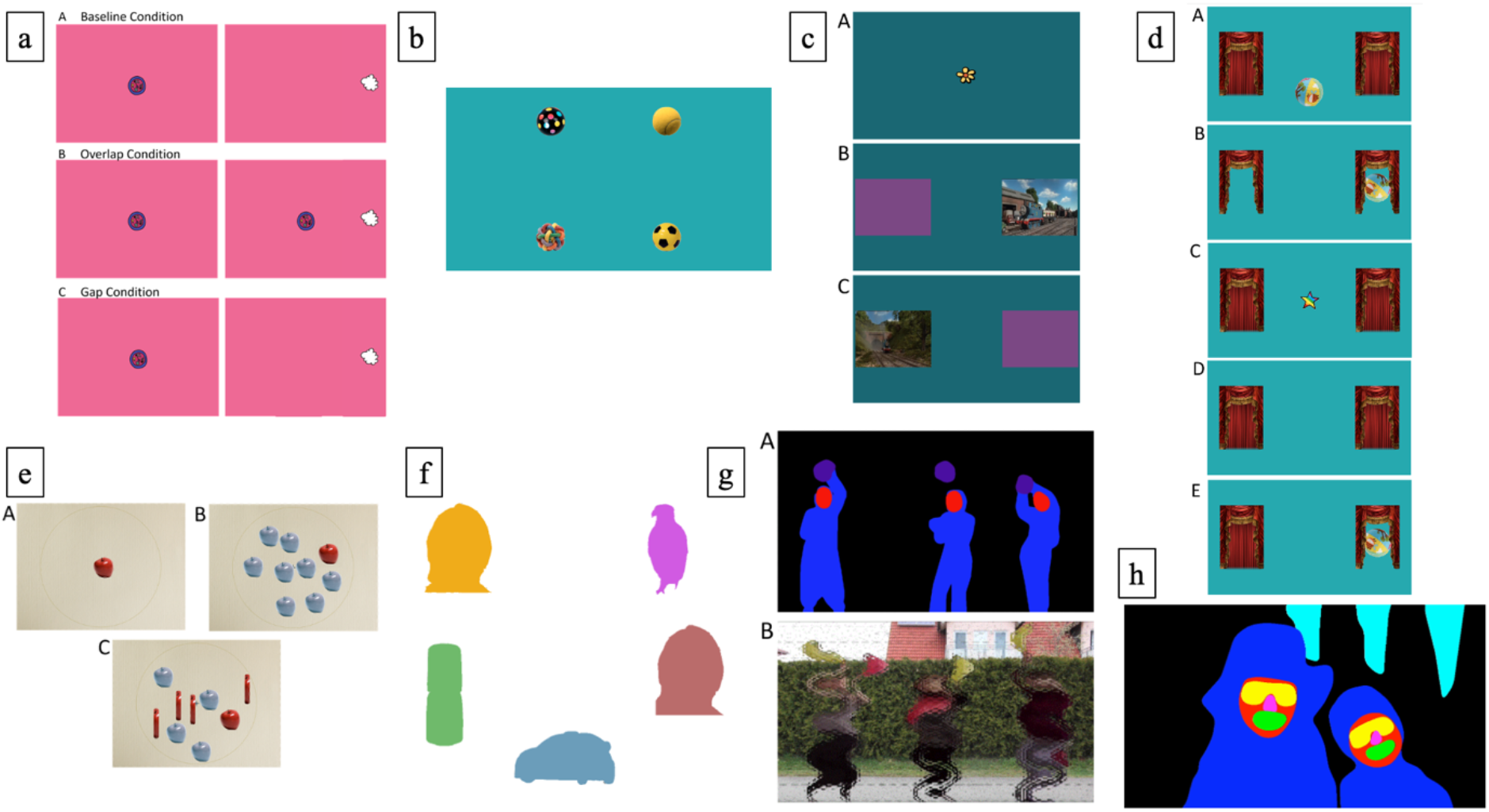
Illustration of the tasks. a) Gap overlap task, showing baseline (A), overlap (B) and gap (C) conditions; b) non-social contingency task; c) reversal learning task, showing the attention-getter (A) and learning (B) and reversal (C) phases; d) working memory task, showing the object appearing (A), moving under a curtain (B), central attention-getter (C), then choice of curtain (D) and revealing of reward (E); e) visual search task pop out task, showing the target (A),single (B) and conjunctive (C) search conditions; f) pop-out task; g) dancing ladies task in both unscrambled (A) and scrambled (B) conditions; h) fifty faces video task.

**Figure 1:**
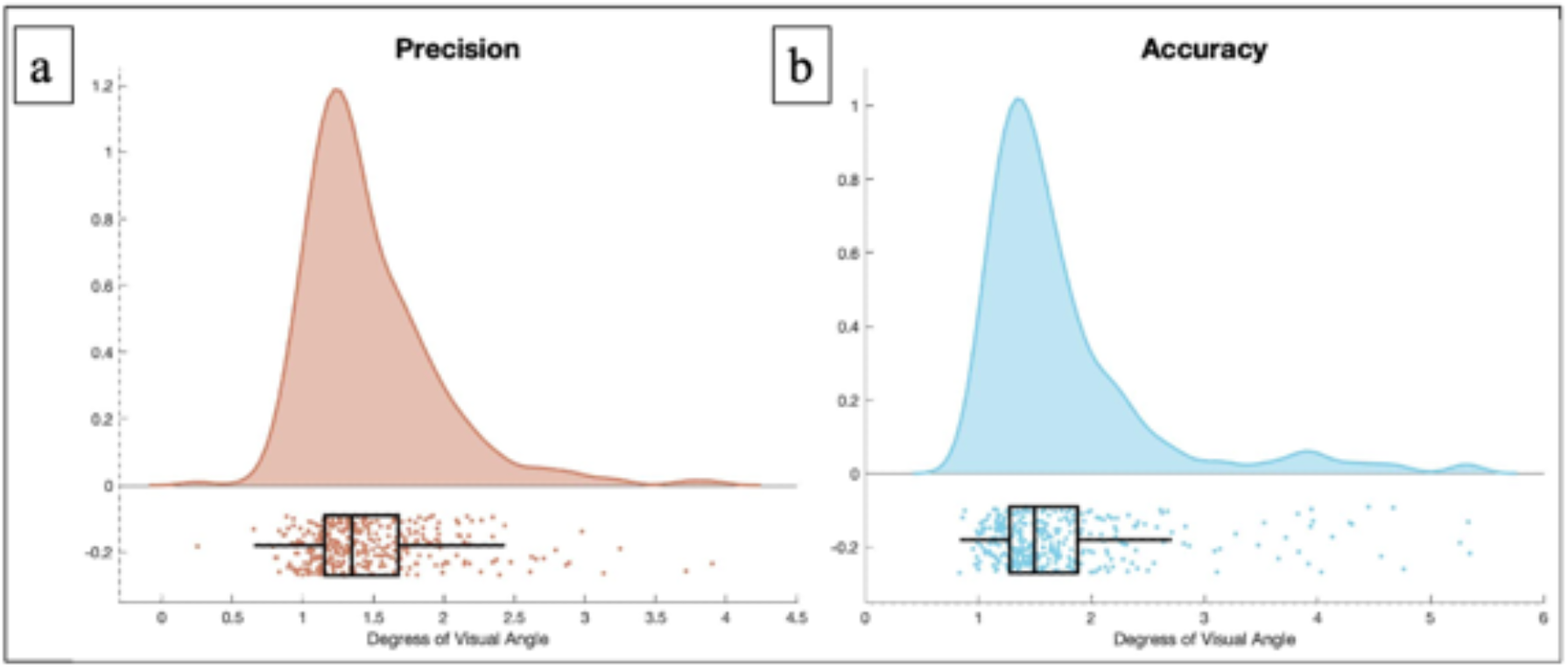
Raincloud plots for (a) precision and (b) accuracy of eye-tracking in the whole sample. *All raincloud plots in this paper were based on Allen et al. (2021)*.

**Figure 3:**
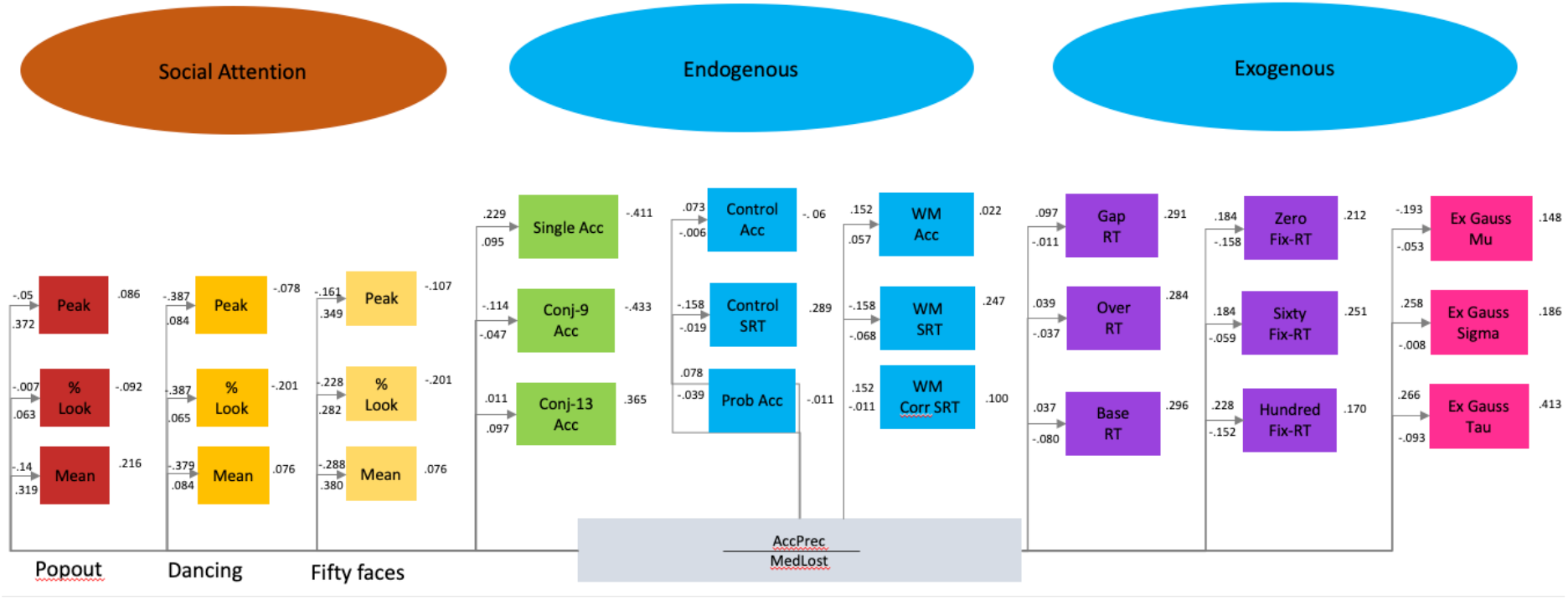
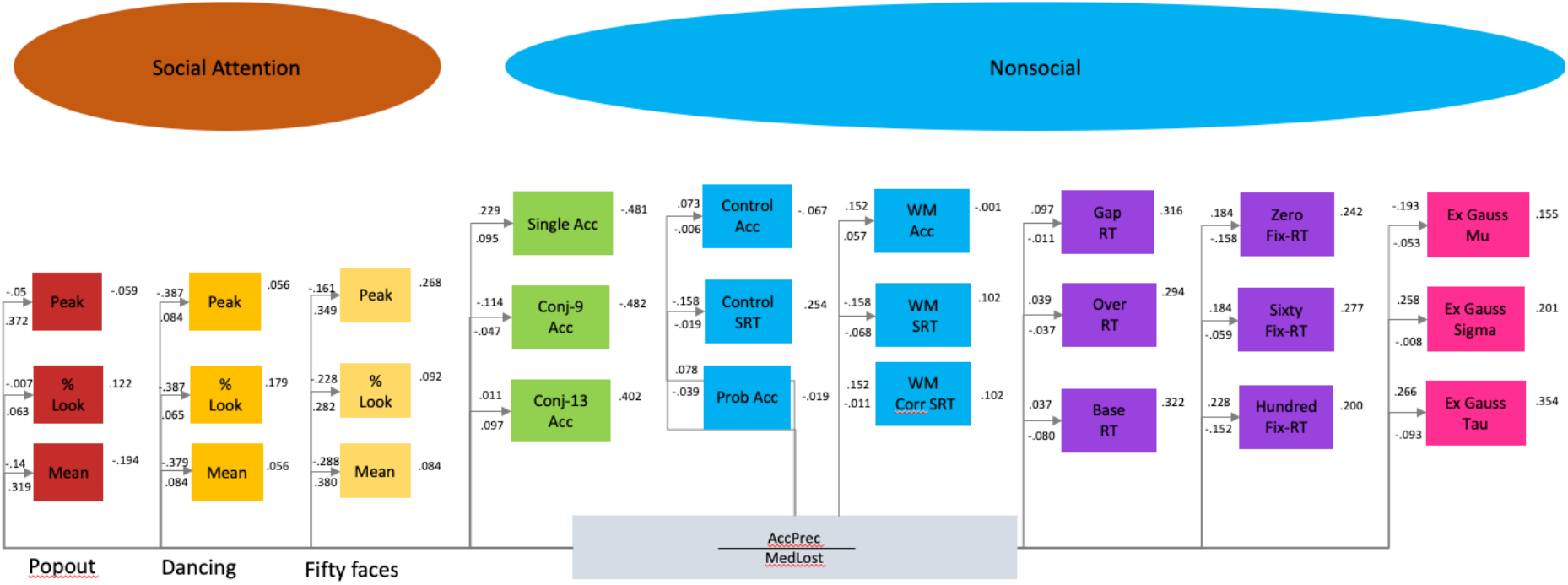
Structual Equation Model of the structure of visual attention within the current eyetracking battery. Although our original model with three factors (a) provided a good fit to the data, it was not significantly better than a model with two latent variables representing social and non-social attention (b).

### 3.3 Condition differences

The expected pattern of condition differences was observed in each task, apart from the working memory task. Briefly, a) in the gap-overlap (SM3.2.2) there were significantly longer reaction times in the overlap than baseline condition, and in the baseline than gap condition (F(1.68, 556.52) = 1015.76, *n* = 332, *p* < 0.001, η_p_^2^ = 0.75’ SM2.2.1); b) in the non-social contingency task (SM3.2.2), times to return to the fixation stimulus in the sixty condition (*M* = 488.06ms, *SD* = 170.78) were faster than in both the zero (*M* = 553.21ms, *SD* = 285.92) and hundred conditions (*M* = 661.82ms, *SD* = 352.26); (Overall condition, F(1.89, 475.86) = 37.88, *p* < 0.001, η_p_^2^ = 0.13; sixty versus zero, F(1, 252) = 11.90, p = 0.001, η_p_^2^ = 0.05; sixty versus hundred, F(1, 252) = 59.69, *p* < 0.001, η_p_^2^ = 0.19 ; SM2.2.2); c) in the reversal learning task (SM3.2.3), accuracy was significantly higher in the pre-switch phase (<u>CC-Pre-Acc)</u> (*M* = 0.83, *SD* = 0.17) compared to the post-switch phase (<u>CC-Post-Acc)</u> (*M* = 0.78, *SD* = 0.25, *n* = 203), *t*(202) = 2.10, *p* = 0.04, CIs 95%[0.003, 0.10], *d* = 0.23); d) in the working memory task (SM2.2.4), a one-sample two-tailed t-test revealed that the proportion of trials in which participants chose the correct location (<u>WM-Acc)</u> was not significantly different to chance, *t*(330) = -1.91, *p* = 0.06, [-0.04, 0.001], *d* = - 0.11, *M* = 0.48, *SD* = 0.19; e) in the visual search task (SM2.2.5), reaction times were slower in the conjunctive (*M* = 1316.68ms, *SD* = 36.12) than single feature displays (*M* = 938.79ms, *SD* = 241.21) with nine elements; (Overall effect of condition in ANOVA, *F*(1.76, 450.90) = 159.90, *p* < 0.001, *n* = 257, η_p_^2^ = 0.38; <u>VS-S9-SRT</u> versus <u>VS-C9-SRT</u>, *F*(1, 256) = 255.46, *p* < 0.001, η_p_^2^ = 0.50) and slower in the conjunctive thirteen (*M* = 1391.81ms, *SD* = 372.28) than conjunctive nine conditions; (F(1, 256) = 5.62, *p* = 0.02, η_p_^2^ = 0.02; Figure 6); f) in the Face pop-out (SM2.2.6), there was more looking to faces than cars and scrambled faces (Overall, *F*(1.39, 354.20) = 165.96, *p* < 0.001, η_p_^2^ = 0.39; face versus car, *F*(1, 255) = 60.31, *p* < 0.001, η_p_^2^ = 0.19; face versus noise, *F*(1, 255) = 607.29, *p* < 0.001, η_p_^2^ = 0.70; means in table 3); g) in the Dancing Ladies videos (SM2.2.7), proportion looking to faces (*M* = 0.14, se = 0.01) was also significantly higher than to objects (*M* = 0.10, se = 0.002), (*F*(1, 248) = 29.66, *p* < 0.001, η_p_^2^ = 0.11), and proportion looking to faces was significantly higher in social (*M* = 0.20, sd = 0.14) than scrambled conditions (*M* = 0.07, *SD* = 0.06), (*t*(248)) = 16.67, *p* < 0.001, *d* = 1.21); h) in the fifty faces video (SM2.2.8) proportion looking to faces (*M* = 0.57, *SD* = 0.15) was also significantly higher than to background people (*M* = 0.03, *SD* = 0.03), (*t*(245) = 53.83, p < 0.001, *d* = 4.99). Thus, each task (with the exception of working memory) replicated previously-reported condition effects or showed the predicted pattern, confirming they can be robustly combined within a large-scale battery.

### 3.3. Factor structure

We first inspected bivariate correlations between core variables from each task (reaction time/accuracy in the cognitive control and working memory; reaction time in the gap; reaction time and proportion of trials retained in the non-social contingency; % looking and peak look to faces in the pop-out, dancing ladies and fifty faces tasks; accuracy in the visual search). This showed strong intercorrelations between variables from the same task, and weaker correlations between tasks. Based on this information and literature-based hypotheses, we conducted a theoretically-motivated structural equation model of the visual attention battery (using sem in Lavaan). We initially tested a three-factor structure with three variables from each of three tasks per construct (proportion looking to faces, peak look and mean look duration to faces from face pop-out, dancing ladies and fifty faces), exogenous orienting (saccadic task reaction times in the three gap-overlap conditions, fixation reaction time in the three conditions of the non-social contingency task, mu, sigma and tau parameters of the exGaussian modelling of reaction times) and endogenous attention (working memory accuracy and reaction time to correct and all trials, cognitive control accuracy in the learning conditions and reaction time during learning, and search accuracy across three conditions in the visual search; Figure 2a). This produced a good fit to the data n=194 RMSEA = 0.034 (CI = 0.019-0.045, p = .99); CFI = 0.975; TLI = 0.9656; AIC = 11551; BIC = 12012; (χ2 (294) = 358, p =0.006, χ ^2^/df=1.21. However, model comparisons showed that a two-factor structure with the exogenous and endogenous factors combined was not a significantly worse fit to the data (χ2 diff(2)= 1.61, p = 0.45) and was more parsimonious; we thus retained the two-factor model, which separated social and non-social attention. Two data quality measures were also included based on a factor analytic decomposition of accuracy and precision and core data quality measures from each task; this revealed two factors reflecting accuracy and precision (21% of the variance) and median lost samples in the free viewing tasks (20% of the variance). The model produced a good fit using both multiple imputation to derive scores from all children (n=350 RMSEA = 0.033 (CI = 0.025-0.040, p = 1); CFI = 0.969; TLI = 0.958; AIC = 19340; BIC = 19988; χ ^2^(296) = 408, p < 0.001, (χ2/df=1.38) and complete cases only (n=194 RMSEA = 0.034 (CI = 0.02-0.046, p = .99); CFI = 0.974; TLI = 0.965; AIC = 11552; BIC = 12006; (χ2 (296) = 363, p =0.005, χ ^2^/df=1.22; (Figure 3). Further, this model was a significant improvement on a model with only one latent factor (n=194 RMSEA = 0.037 (CI = 0.024-0.048, p = .98); CFI = 0.970; TLI = 0.959; AIC = 11563; BIC = 12014; (χ2 (297) = 376, p = 0.001, χ ^2^/df=1.27; χ2 diff(3)= 13.13, p = 0.0003). The two latent variables were not associated (n=194 B=0.087 SE=0.084 z=1.04, p = 0.3). Higher scores for social attention represent more interest in social content; higher scores for non-social attention represent slower and less accurate cognitive responses (Figure 2b). These summary scores may prove useful for investigators wishing to represent distinct sources of variance in the battery as a whole.

## Discussion

Although much existing research has used eye-tracking to assess visual attention in infants and young children, little work has considered the feasibility of conducting large-scale eye-tracking studies in toddlers. In this study, we collected data from a battery of eight eye-tracking tasks completed by over three hundred 18-month-olds. Data were successfully collected from 99.1% (347) of toddlers on at least one task. Analyses found expected condition effects in seven out of eight eye-tracking tasks, with only the working memory task not finding the expected effects at the group level. A hypothesis-driven SEM provided a good fit to the data, indicating that in addition to indices from individual tasks, the battery can be used to extract global measures of social and non-social attention. Using maximum likelihood imputation allows latent variable measures to be extracted for all infants in the sample. Overall, the current eye-tracking battery provides a feasible and objective measure of visual attention in 18-month-old children.

### Overall feasibility

Overall feasibility of the study was good. For every task at least 70% of children provided enough data to be included in analyses; this was 90% for four of the first five tasks in the battery. After cut-off criteria were applied, at least 64% of the whole sample were retained; over 86% for the four tasks with highest retention. Fewer than 10% of children failed to meet cut-off criteria, indicating that the majority of participants who completed the task produced valid data and supporting the feasibility of the eye-tracking tasks in this age range. Furthermore, all but one condition effects were the same if children who did and did not meet typical cut-offs were included, indicating that patterns of performance reported in this age range do not just capture effects seen in children with stronger attentiveness.

Despite validation procedures, data quality still impacted extracted metrics. For free-viewing tasks, data duration was associated with looking to faces, and the degree to which looking to faces was greater than comparison stimuli. One potential explanation is that children who spend less time looking at the stimuli have less time to process the content and show less differential endogenously-mediated attention to stimulus features. Alternatively, it may be that children who are less interested in faces and people spend less time looking at the videos, which feature prominent social content. Such considerations are important to balance when interpreting relations between variables of interest and data quality. Metrics which were associated with either accuracy or precision metrics at a level of >0.2 (suggesting they should be controlled in research using these tasks) include gap reaction time in the Gap-Overlap task, reaction times in the non-social contingency task, singleton nine visual search accuracy, percentage looking to faces and multiple peak look duration variables in the dancing ladies and percentage looking and peak look duration to faces in the Fifty Faces free viewing video. This is likely because even with expanded areas of interest, lower accuracy and precision leads to less certainty in gaze sample classification. Loadings in the model suggested precision and accuracy measures may be better captured as one data quality factor, with median lost samples another; as such, controlling for these factors may be the most parsimonious approach.

### Task robustness

The majority of tasks in the current battery produced robust condition effects in the expected direction. These included competition effects in visual orienting, the ability to use memory to find a video, set size and pop-out effects on search for a visual target and the preference for faces in early development. These findings indicate that it is possible to robustly measure many previously observed effects, even within a large battery of tasks. This has important implications for the replicability and validity of these findings, strengthening support for their generalisability and robustness. In the light of increasing concerns about the reproducibility of cognitive psychology (Huber et al., 2019), this set of replications provides important reassurance that developmental science has produced a range of robust observations about the developing infant attention system.

The one exception was the working memory task, which only provided some weak evidence of working memory at the group level when trial-level reaction time data were analysed. This may reflect the fact that this skill is relatively fragile at this age; indeed, there are few robust demonstrations of single-trial level working memory success at 18 months (Hendry et al., 2016). One important task feature may be that the child had to actively ‘find’ the object with their gaze, rather than passively watch sequences of objects as has previously been used in other working memory tasks with younger infants (Ahmed & Ruffman, 1998; Baillargeon et al., 1985). Indeed, Hood et al. (2003) found that the same two-and-a-half-year-olds demonstrated ability to recognise impossible locations of an object but failed to complete active search and retrieval of the object, indicating a disparity between active search and passive observation. Although children typically succeed at passive visual tasks at a substantially younger age than behavioural analogues (Ahmed & Ruffman, 1998; Baillargeon & Graber, 1988; Hofstadter & Reznick, 1996), it may be that this active component makes tasks more difficult.

### Structure of visual attention

Once strong within-task associations were accounted for, a hypothesis-driven SEM provided a good fit to the data. The most parsimonious structure indicated two latent variables that can be interpreted as social attention (interest in people) and non-social attention (speed and accuracy of saccades). We used overall measures of accuracy and reaction time in this model because condition difference scores can be unstable (combining the noise inherent in both variables). Indeed, Draheim et al. (2019) suggested that the lack of correlation between attention measures in many studies may be due to methodological issues with attention capture tasks, rather than that attention not representing an unified concept.

They conclude that accuracy-based measures may provide more reliable and valid measures of attention control than reaction times and difference scores. This assessment was based on a number of judgements, including how well metrics from various tasks inter-correlated with one another; notably, considerable correlations were found even between metrics from tasks which make markedly different demands on the participant.

The lack of correlations between the two latent variables in our SEM is consistent with theoretical models in which social attention represents a distinct construct. The large sample size in the current work implies this result is not due to lack of statistical power. Based on this, eye-tracking metrics should be carefully chosen in future work in this field, to ensure that tasks measure an appropriate facet of visual attention. Although many models of attention distinguish exogenous and endogenous orienting, inclusion of this distinction in our modelling did not explain significantly more variance than a two-factor model. Neuroimaging studies have indicated that both endogenous and exogenous attention shifts are mediated by the same large-scale fronto-parietal networks, indicating they may be closely related constructs (Peelen et al., 2004). However, other studies have shown distinct impacts of environmental variation (such as screen exposure) on exogenous and endogenous attention shifting (Portugal et al., 2021). A model incorporating the endogenous/exogenous distinction did provide an adequate fit to the data, and may be preferred by investigators who wish to distinguish these components of visual attention at the individual level.

Though this work draws strengths from its large sample size and substantial task battery, there are some limitations. Whilst it is assumed that eye-tracking tasks provide a direct measure of attention, assumptions are made about specifically what aspect of attention tasks are tapping. Most tasks in the current battery have been widely used previously; the current study serves to establish previously-found condition effects in this large toddler sample. Yet, inaccurate assumptions about the processes captured by different tasks may lead to mistakes in how effects are interpreted. Additionally, processing of data has been carried out in line with other work using these tasks, but this could be problematic. Measures calculated directly from the eye-tracker remove an element of subjective assessment, but there remain numerous options for processing; some of these decisions may impact the results found in later analyses. Such limitations ought to be considered by eye-tracking researchers. Indeed, there is a continuing effort to establish reliable and valid eye-tracking batteries in young children (van Baar et al., 2020); the current work contributes to this field by establishing the robustness of these eye-tracking tasks.

The current study demonstrates that this large eye-tracking battery can be successfully used with 18-month-olds and produces both task-based and cross-task indices that may be useful for large-scale assessment of visual attention in toddlers. Our dataset is embedded in a large-scale study of child development (http://www.developingconnectome.org/) that also includes measures of neonatal brain structure (Bozek et al., 2018) and function (Eyre et al., 2021) and clinical and developmental assessments in toddlerhood. This context provides a strong foundation for investigators to examine the relation between individual differences in attentional processing and other aspects of child development. Further, investigators can access our task battery (built in TaskEngine; https://sites.google.com/site/taskenginedoc/) for use in their own large-scale cohort studies, facilitating greater robustness and comparability in large scale studies of child development; ongoing efforts include assessments in infants with a family history of autism (Jones et al., 2019), infants with a history of heart problems and premature infants. In summary, we provide a resource for investigators to further probe the role of individual differences in visual attention in toddlerhood, and their relation to structural brain development and clinical outcomes.

## Acknowledgments

We are grateful for all the families who kindly agreed to participate in the project and recognize their particular commitment in remaining engaged with the programme during the COVID-19 Pandemic. We also acknowledge the support of the Neonatal Intensive Care Unit and the Newborn Imaging Centre at Evelina London Children’s Hospital. We also thank the external advisory board for their expert advice and contribution across the dHCP project: David van Essen, Arthur Toga, Richard Frackowiak, Dan Marcus, Petra Huppi, Essa Yacoub, and John Ashburner.

