## Supplementary Materials for "Objective assessment of visual attention in toddlerhood"

### Objective assessment of visual attention in toddlerhood: Supplementary Materials

#### 1. Additional Methods

##### 1.1 Eye tracking data acquisition

Raw eye tracking data was acquired via the Tobii Gaze Analytics SDK 3.0, processed and saved to disk. Trial onset and offset was associated with the current sample of gaze data, and time-stamped in the eye tracker's time format. When a video was playing, an additional timestamp was recorded every 30 frames, in order to ensure constant synchronisation between stimuli and data. The screen of the TX-300 has a diagonal size of 23" (58.42cm x 28.6cm, 52.0° x 26.8° @ 60cm), a native resolution of 1920 x 1080 pixels and an aspect ratio of 16:9. Eye-tracking variables are named to match the data release available via the Developing Human Connectome Project (DHCP) (http://www.developingconnectome.org/project/).

##### 1.2 Tasks

If the participant became bored or fussy, the experimenter could skip the current trial and move on to the next. Skipped trials were marked in the data and excluded from analysis. Blocks of each task were presented interspersed with each other to maintain child attention in a pseudorandomised order. Eye-tracking variables are named to match the data release, available via the Developing Human Connectome Project (DHCP) (http://www.developingconnectome.org/project/).

###### *1.2.1 Gap-Overlap*

*1.2.1.1. Stimulus presentation*

All stimuli were presented at a size of 3cm x 3cm (2.86° x 2.86° at 60cm viewing distance). Reward stimuli were either a star, a sun, a dog, cat, pig, tiger or tortoise which were animated and accompanied by a sound.

*1.2.1.2. Data extraction*

Data were analysed offline. Each trial was inspected automatically to determine trial validity and calculate a saccadic reaction time (SRT) to shift attention from the CS to the PS, relative to PS onset. A trial was valid if the following conditions were met: 1) gaze fell on the CS; 2) no gaps of missing data longer than 200ms were present during the CS period (before PS onset); 3) there was at least one sample of gaze on the CS within 50ms either side of PS onset; 4) no gaps of missing data longer than 100ms were present during the PS period (between PS onset and reward onset); 5) SRT was longer than 150ms and shorter than 1200ms; 6) gaze did not go in the opposite direction to the side of the PS; 7) gaze did not enter the PS AOI after engagement with the CS but before PS onset.

###### *1.2.2 Reversal learning ('Cognitive Control')*

*1.2.2.1. Stimulus presentation*

Participants were presented with two blocks of 18 trials. Rectangles (17cm x 12.5cm, 16.1° x 11.9° @ 60cm) were 0.5cm (0.48°) from each edge of the screen and vertically-centred.

*1.2.2.2. Data extraction*

Data were analysed offline. The first trial of each phase, where the participant was not yet aware of the location of the video, was discarded. Pre-switch and post-switch anticipation (CC-Pre-Ant and CC-Post-Ant), calculated as *number of trials in which participants fixated a rectangle (whether it was correct or not) / total number of valid trials* within the pre-switch and post-switch phases, were included as control variables.

##### *1.2.3 Working memory*

###### *1.2.3.1. Stimulus presentation*

Participants were presented with three blocks of five trials. Theatre stage images were 16cm x 11cm, 15.2° x 10.5° and were presented 3cm (2.9°) from the edge. Trials began with the appearance of a horizontally-centred child's toy, selected randomly from 19 exemplars. The toy was 7cm (6.9°) wide, and the height was adjusted to match the aspect ratio of each individual image.

##### *1.2.4 Visual search*

###### *1.2.4.1. Stimulus presentation*

To highlight the special status of the target through pop-out, the first three test trials consisted of single feature displays. To emphasize this further and to grab participant's attention and fixation, before each trial began, the target (a red apple) 'flew in' from the upper portion of the screen, stopped in the centre of the screen for one second then disappeared. Aside from these first three, displays within each test trial were mixed in blocks and presented in random order. In all trials, a sound effect accompanied each event visual event.

##### *1.2.5 Face Pop-out*

###### *1.2.5.1. Stimulus presentation*

Each array was presented for 10 seconds and counter-balanced for the location of the face in the array. The stimulus array was presented full-screen with adjustments for a proper aspect ratio, at 43.8cm x 28.6cm (39.0° x 26.8° @ 60cm).

###### *1.2.5.2. Data extraction*

Each AOI was scored by counting the number of samples of gaze data that fell on each AOI. Trials were marked as invalid if either a) the proportion of valid (non-missing) samples was less than 25%, or b) the duration of data was less than 5s.

#### **1.3. Data quality assessment**

Accuracy and precision were calculated during the gaze-contingent fixation stimulus that preceded each trial. The AOI around the fixation stimulus was 1.75x larger than the stimulus itself. The trial would begin even under conditions of high accuracy drift. Because the fixation stimulus was always at a known location, and because the trial would not begin until that location was fixated, we can use it to calculate the spatial error between the true gaze location and the gaze location reported by the eye tracker. The AOI around the fixation stimulus was 1.75x larger than the stimulus itself. The trial would begin even under conditions of high accuracy drift. Accuracy was calculated as the root-mean-square (RMS) of the euclidean distance between the location of each gaze sample and the location of the fixation stimulus. Precision was calculated as the RMS of the euclidean distance between each gaze sample and the centroid of all gaze samples.

**1.4. Analytic strategy**

Expected condition effects were: (i) Gap-Overlap: longer reaction times for the Overlap than Baseline and Baseline than Gap condition (Elsabbagh et al., 2009, 2013); (ii) Popout: more looking at the face and a greater proportion of trials on which the first look was to the face (Gliga et al., 2009); (iii) Reversal learning: greater accuracy on the learning than reversal trials; (iv) Working memory: detection of the correct location significantly more than chance; (v) Nonsocial contingency: more attention to the last static slide and faster reaction times to select a ball during the trials in the 60% vs 100% and 0% conditions; (v) Dancing ladies: longer looking to faces than objects or background; (vii) 50 faces: longer looking to faces than objects, and show stronger effects for the native than degraded conditions; (viii) Visual search: faster reaction times and greater accuracy during the conjunctive nine condition. For all analyses, where sphericity was not met a Greenhouse-Geisser was applied and where Levene's test was significant, equal variances were not assumed. All data presented in this study will be made available via the Developing Human Connectome Project (DHCP) open access data release (http://www.developingconnectome.org/project/).

#### 2. Additional results

**2.1. Data Quality analyses**

*2.1.1. General eye-tracking data quality*

Two measures of the general quality of the eye-tracking data across the session were calculated. Accuracy (the spatial displacement of recorded gaze from the point fixated) and precision (variability in consecutive samples on the same fixation point) were extracted as proxies of general eye-tracking quality across the session. Table S3 shows summary statistics of accuracy and precision data for the sample of participants who provided data for any task and for whom data quality data could be extracted. Table S4 shows associations between accuracy and precision and each of the key task variables; these associations include participants who provided enough data to be included in condition effect analyses for each task before cut-off criteria were applied.

Table S4: *Mean, standard deviation, range and number for precision and accuracy measures*

| | *Mean* | *SD* | *Range* *Minimum* | *Maximum* | *n* |
|---|---|---|---|---|---|
| Accuracy | 1.74 | 0.79 | 0.83 | 5.35 | 340 |
| Precision | 1.47 | 0.47 | 0.25 | 3.90 | 340 |

Table S5: *Associations between each of accuracy and precision, and key task variables*

| Task | Variable | Accuracy | | Precision | | |
|---|---|---|---|---|---|---|
| | | *r*-value | *p*-value | *r*-value | *p*-value | *n* |

| | | | | | | |
|---|---|---|---|---|---|---|
| Gap | GO-Gap-SRT | .25 | < 0.001 | 0.18 | 0.001 | 333 |
| | GO-Baseline-SRT | .11 | 0.04 | .02 | 0.74 | 332 |
| | GO-Overlap-SRT | .05 | 0.36 | .03 | 0.61 | 333 |
| | GO-Facilitation | -.15 | 0.007 | -.19 | 0.001 | 331 |
| | GO-Disengagement | -.01 | 0.83 | 0.01 | 0.82 | 331 |
| Non-social contingency | NSC-100-SRT-NoZone | -.06 | 0.36 | -.07 | 0.27 | 282 |
| | NSC-60-SRT-NoZone | -.09 | 0.11 | -.10 | 0.08 | 323 |
| | NSC-0-SRT-NoZone | -.25 | < 0.001 | -.20 | 0.001 | 271 |
| | NSC-100-FixRT | .26 | < 0.001 | .22 | < 0.001 | 282 |
| | NSC-60-FixRT | .22 | < 0.001 | .23 | < 0.001 | 324 |
| | NSC-0-FixRT | .19 | 0.002 | .27 | < 0.001 | 271 |
| Reversal Learning | RL-Pre-Acc | .06 | 0.32 | .14 | 0.01 | 320 |
| | RL-Post-Acc | -.05 | 0.48 | -.10 | 0.15 | 203 |
| | RL-Pre-SRT | -.05 | 0.40 | -.08 | 0.16 | 320 |
| | RL-Post-SRT | .03 | 0.65 | -.08 | 0.27 | 203 |
| Working Memory | WM-Acc | .05 | 0.39 | .11 | 0.04 | 330 |
| | WM-SRT-Acc | -.05 | 0.33 | -.13 | 0.02 | 323 |
| | WM-SRT-Inacc | -.05 | 0.38 | .13 | 0.02 | 327 |
| | WM-SRT | -.04 | 0.52 | -.15 | 0.005 | 330 |
| Visual Search | VS-S9-SRT | .06 | 0.31 | .01 | 0.90 | 259 |
| | VS-C9-SRT | -.07 | 0.27 | -.12 | 0.05 | 257 |
| | VS-C13-SRT | -.07 | 0.30 | -.11 | 0.07 | 257 |
| | VS-S9-Acc | -.32 | < 0.001 | -.16 | 0.003 | 336 |
| | VS-C9-Acc | -.15 | 0.005 | -.01 | 0.86 | 336 |
| | VS-C13-Acc | -.09 | 0.12 | .03 | 0.61 | 336 |
| Pop-out | Pop-Face-Pct | -.10 | 0.12 | -.10 | 0.12 | 256 |
| | Pop-Car-Pct | -.11 | 0.09 | -.06 | 0.32 | 256 |
| | Pop-Noise-Pct | -.09 | 0.17 | -.07 | 0.30 | 256 |
| | Pop-Face-Peak | -.12 | 0.06 | -.16 | 0.01 | 255 |

| | | | | | | |
|---|---|---|---|---|---|---|
| | Pop-Car-Peak | -.14 | 0.03 | -.12 | 0.05 | 254 |
| | Pop-Noise-Peak | -.05 | 0.46 | -.11 | 0.07 | 252 |
| Dancing Ladies | Dance-Soc-Face-Pct | -.39 | < 0.001 | -.29 | < 0.001 | 250 |
| | Dance-Soc-Object-Pct | .05 | 0.45 | -.05 | 0.45 | 250 |
| | Dance-Scr-Face-Pct | -.13 | 0.04 | -.05 | 0.44 | 250 |
| | Dance-Scr-Object-Pct | -.17 | 0.006 | -.18 | 0.004 | 250 |
| | Dance-Soc-Face-Peak | -.37 | < 0.001 | -.30 | < 0.001 | 249 |
| | Dance-Soc-Object-Peak | .08 | 0.19 | -.07 | 0.30 | 250 |
| | Dance-Scr-Face-Peak | -.20 | 0.001 | -.17 | 0.007 | 244 |
| | Dance-Scr-Object-Peak | -.17 | 0.009 | -.22 | 0.001 | 249 |
| Fifty Faces | 50Face-Face-Peak | -.21 | 0.001 | -.23 | < 0.001 | 244 |
| | 50Face-Background-Peak | .05 | 0.43 | -.02 | 0.78 | 228 |
| | 50Face-Face-Pct | -.28 | < 0.001 | -.19 | 0.003 | 246 |
| | 50Face-Background-Pct | .04 | 0.50 | .003 | 0.96 | 246 |

#### 2.2. Task-specific analyses

*2.2.1 Gap-Overlap task*

94.9% of children successfully completed the gap task with an average of 13 valid trials per condition. A one-way repeated-measures ANOVA comparing reaction times in gap, overlap and baseline conditions indicated reaction times significantly varied with condition, ($F(1.68, 556.52) = 1015.76$, $n = 332$, $p < 0.001$, $\eta_p^2 = 0.75$); simple comparisons indicated (as expected) longer reaction times for the overlap ($M = 590.45$ms, $SD = 18.18$) than baseline ($M = 575.11$ms, $SD = 13.45$), ($F(1, 331) = 1056.474$, $p < 0.001$, $\eta_p^2 = 0.76$), and baseline than gap ($M = 555.30$ms, $SD = 13.33$), ($F(1, 331) = 372.501$, $p < 0.001$, $\eta_p^2 = 0.53$; Figure 2). Similar patterns were seen when restricting analysis to children with >5 trials/condition (SM*2.3.2.1.*) or accounting for trial variability (SM*2.2.2.1.*). Thus, this task yielded good data quantity and quality and the expected condition effects when administered in a longer battery.

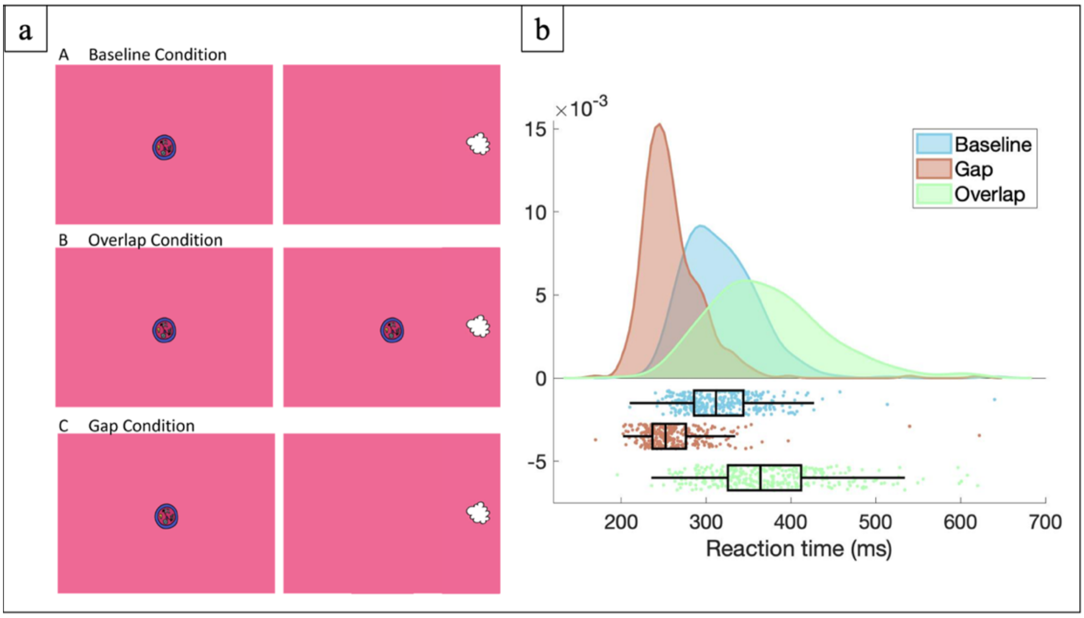


*Figure 2:* (a) Gap-overlap task display and (b) raincloud plot of reaction times in gap, overlap and baseline conditions of the gap-overlap task.

*2.2.2 Non-social Contingency*

72.3% of children successfully completed all three conditions with an average of 19 valid trials per condition. Multiple one-way repeated measures ANOVAs were conducted to compare two reaction time variables across three conditions (zero, sixty and hundred); Figure 3. Reaction time to select a ball did not differ between the sixty ($M$ = 537.95ms, $SD$ = 53.95) and hundred ($M$ = 536.53ms, $SD$ = 76.48), or between the zero and hundred conditions but, was significantly longer in the sixty versus zero condition as expected ($M$ = 526.33ms, $SD$ = 67.75); (Overall condition, $F(1.89, 473.55) = 2.93$, p = 0.06, $\eta_p^2 = 0.01$; sixty versus hundred, F(1, 251) = 0.07, p = 0.80, $\eta_p^2 = 0.00$; zero versus sixty, $F(2, 251) = 6.52$, $p = 0.01$, $\eta_p^2 = 0.03$; zero versus hundred, $F(1, 251) = 3.43$, $p = 0.07$, $\eta_p^2 = 0.01$). For times to return to the fixation stimulus, reaction times in the sixty condition ($M$ = 488.06ms, $SD$ = 170.78) were faster than in both the zero ($M$ = 553.21ms, $SD$ = 285.92) and hundred conditions ($M$ = 661.82ms, $SD$ = 352.26); (Overall condition, F(1.89, 475.86) = 37.88, $p < 0.001$, $\eta_p^2 = 0.13$; sixty versus zero, F(1, 252) = 11.90, p = 0.001, $\eta_p^2 = 0.05$; sixty versus hundred, F(1, 252) = 59.69, $p < 0.001$, $\eta_p^2 = 0.19$).

All children who completed the task met the cut-off criteria, therefore results were identical if children were only included with a minimum of 5 valid trials per condition (SM*2.3.2.2.*). Both reaction time to select a ball and reaction times to return to the fixation stimulus were significantly faster in the sixty versus zero and hundred conditions when analyses were run on trial-level data (SM*2.2.2.2.*). Thus, this task yielded reasonable quantity and good quality of data and expected condition effects were found when the task was administered in a longer battery; accommodating trial-level data may be the more powerful analytical approach.

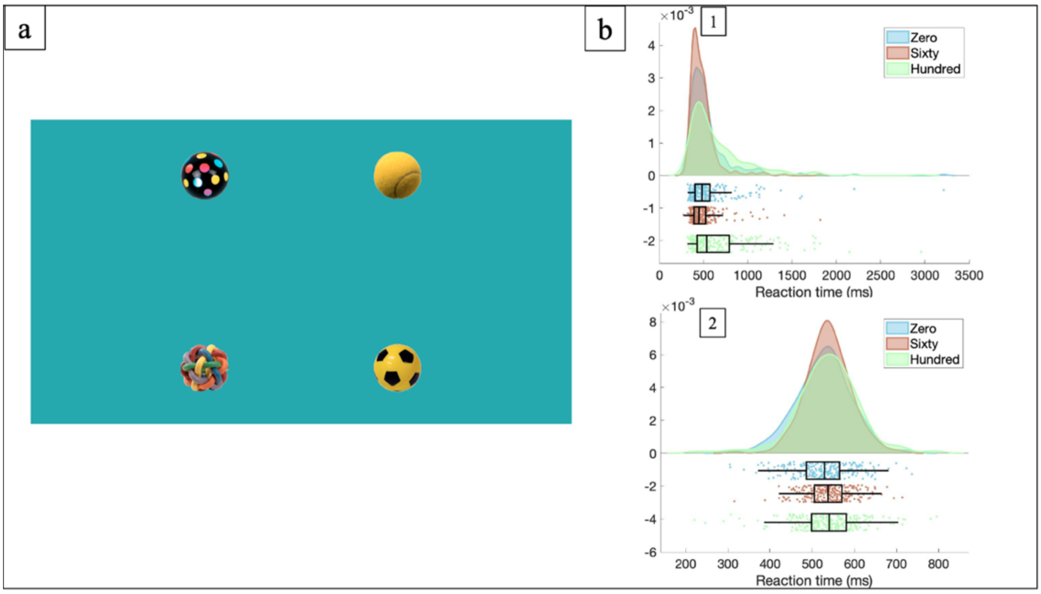


*Figure 3:* (a) Stimulus presentation and (b) raincloud plot of (b1) reaction time with zone-outs removed and (b2) reaction time to return to a central fixation stimulus after a choice in zero, sixty and hundred conditions of the non-social contingency task.

*2.2.3 Reversal learning ('Cognitive Control')*

92.3% of children successfully completed the task with an average of 6 (*SD* = 2) valid trials for the pre-switch condition. During the pre-switch phase of the cognitive control task, the mean proportion of trials in which participants correctly anticipated animation (CC-Pre-Acc) was approximately 70% (*M* = 0.70, *SD* = 0.27, *n* = 322). The mean reaction time for participants to select an AOI (CC-Pre-SRT) was 688.18ms (*SD* = 206.23, *n* = 322). Analyses were also conducted with a control variable, percentage of trials anticipated at all (SM2.1.1.).

Of the group of 323 participants who completed the learning phase of the cognitive control task, 203 (62.8%) participants 'passed' and proceeded to the reversal condition, with an average of 5 (2) valid trials for this condition. Repeated-measures t-tests showed no significant differences in reaction times during the pre-switch (CC-Pre-SRT) (*M* = 653.70ms, *SD* = 183.42) and post-switch phases (CC-Post-SRT) (*M* = 670.61ms, *SD* = 205.53, *n* = 203), *t*(202) = -1.08, *p* = 0.28, CIs 95%[-0.05, 0.01], *d* = 0.10), however accuracy was significantly higher in the pre-switch phase (CC-Pre-Acc) (*M* = 0.83, *SD* = 0.17) compared to the post-switch phase (CC-Post-Acc) (*M* = 0.78, *SD* = 0.25, *n* = 203), *t*(202) = 2.10, *p* = 0.04, CIs 95%[0.003, 0.10], *d* = 0.23, Figure 4. Thus, children who completed both phases of the task showed some reduced performance after reversal, though not in all variables.

Independent t-tests conducted between the two groups who did and did not do the reversal phase found that as expected given the design, a significantly lower proportion of trials were correctly anticipated (CC-Pre-Acc) by the group who did not go on to do the reversal phase; this same group also showed significantly slower reaction times; table S1 & S2.

Results were consistent if children were only included with a minimum of 2 valid trials per condition (SM*2.3.2.3.*) whilst the reaction time effect between pre- and post-switch phase was significant when accounting for trial variability SM*2.2.2.3.*). Given not all children completed the reversal phase and data was not missing at random, further modelling thus used the accuracy and reaction time from the learning phase only. Overall, this task yielded good quantity and quality of data and some evidence to support the expected condition effects when administered in a longer battery.

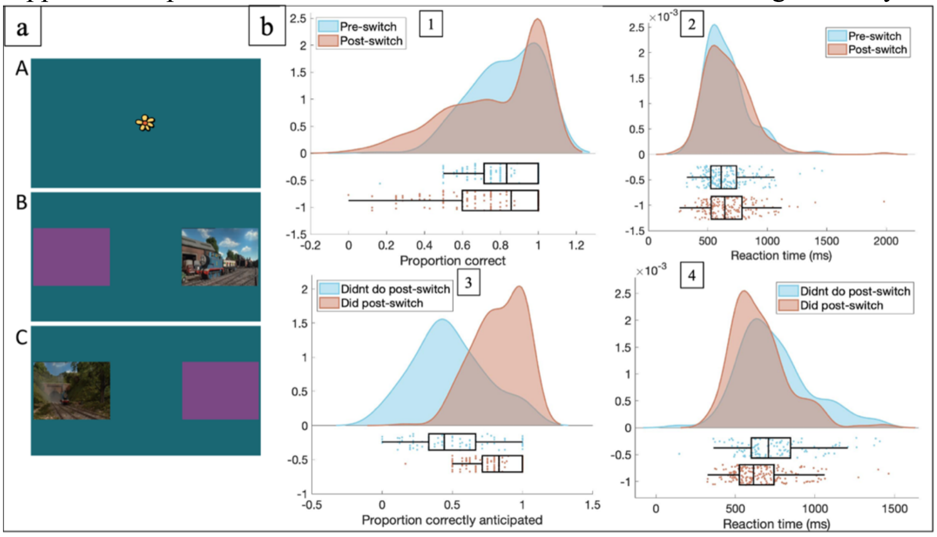


*Figure 4:* (a) Reversal Learning task display and (b) raincloud plot of (b1) the proportion of trials correctly anticipated and (b2) reaction times during the pre- and post-switch phases of the cognitive control task, (b3) the proportion of trials correctly anticipated and (b4) reaction time in the pre-switch phase by groups who did and didn't complete the post-switch phase.

Percentage of trials anticipated (either correctly or incorrectly) was included as a control variable. During the pre-switch phase, the mean proportion of trials in which participants anticipated an animation at all (CC-Pre-Ant) was 89% ($n$ =323, $M$ = 0.89, $SD$ = 0.19).
A repeated t-test found that the proportion of trials anticipated in the pre-switch phase (CC-Pre-Ant); $M$ = 0.96, $SD$ = 0.10; was higher than in the post-switch phase (CC-Post-Ant); $M$ = 0.93, $SD$ = 0.16, t(202) = 2.05, p = 0.04, d = 0.15, CIs 95%[0.0001, 0.05]. An independent t-test conducted between the two groups who did and didn't do the reversal phase found that a significantly lower proportion of trials were anticipated by the group who didn't go on to do the reversal phase, t(142.25) = 7.28, p < 0.001, d = 1.01, CIs 95%[0.12, 0.22], table S2.

Table S1: *t-value, degrees of freedom, p-value, Cohen's d, lower and upper CIs from independent t-tests between the proportion of trials correctly anticipated, anticipated at all and reaction times between two groups who did and didn't do the reversal phase in the cognitive control task*

| | *t* | *df* | *p* | *d* | *lower* | *upper* |
|---|---|---|---|---|---|---|
| CC-Pre-Acc | 13.07 | 179.56 | < 0.001 | 1.67 | 0.29 | 0.40 |
| CC-Pre-SRT | -3.79 | 206.20 | < 0.001 | -2.34 | 44.71 | 141.90 |

Table S2: *Mean, standard deviation and n for groups who did and didn't do the post-switch phase of the cognitive control task*

| Variable | Group | *Mean* | *SD* | *n* |
|---|---|---|---|---|
| Proportion of trials correctly anticipated (CC-Pre-SRT) | Did reversal condition | 0.83 | 0.17 | 203 |
| | Didn't do reversal condition | 0.49 | 0.26 | 119 |
| Reaction time (CC-Pre-SRT) | Did reversal condition | 653.70 | 183.43 | 203 |
| | Didn't do reversal condition | 747.00 | 229.28 | 119 |

| | | | | |
|---|---|---|---|---|
| Proportion of trials anticipated (CC-Pre-Ant) | Did reversal condition | 0.96 | 0.10 | 203 |
| | Didn't do reversal condition | 0.79 | 0.25 | 120 |

*2.2.4 Working Memory*

94.6% of children successfully completed the task with an average of 14 (3) valid trials. A one-sample two-tailed t-test revealed that the proportion of trials in which participants chose the correct location (WM-Acc) was not significantly different to chance, $t(330) = -1.91$, $p = 0.06$, [-0.04, 0.001], $d = -0.11$, $M = 0.48$, $SD = 0.19$.

A paired samples t-test showed that there was no difference in mean reaction times for trials in which participants were correct (WM-SRT-Acc) ($M = 715.9$ms, $SD = 207.4$, $n = 321$) versus incorrect (WM-SRT-Inacc) ($M = 706.7$ms, $SD = 195.4$, $n = 321$), $t(320) = 0.76$, $p = 0.45$, [-14.6, 33.13], $d = 0.05$, Figure 5). Mean reaction time across all trials (WM-SRT-All) was 700.8s ($SD = 182.9$, $n = 324$). Results were consistent if only children with a minimum of 10 valid trials were included (SM*2.3.2.4.*) When analysing at the individual trial level, reaction times were significantly longer in incorrect versus correct trials (SM*2.2.2.4.*), indicating that this may be a more powerful analysis for detecting memory effects. Thus, children did not show evidence of successfully remembering the location of the object on the group level, although there were considerable individual differences.

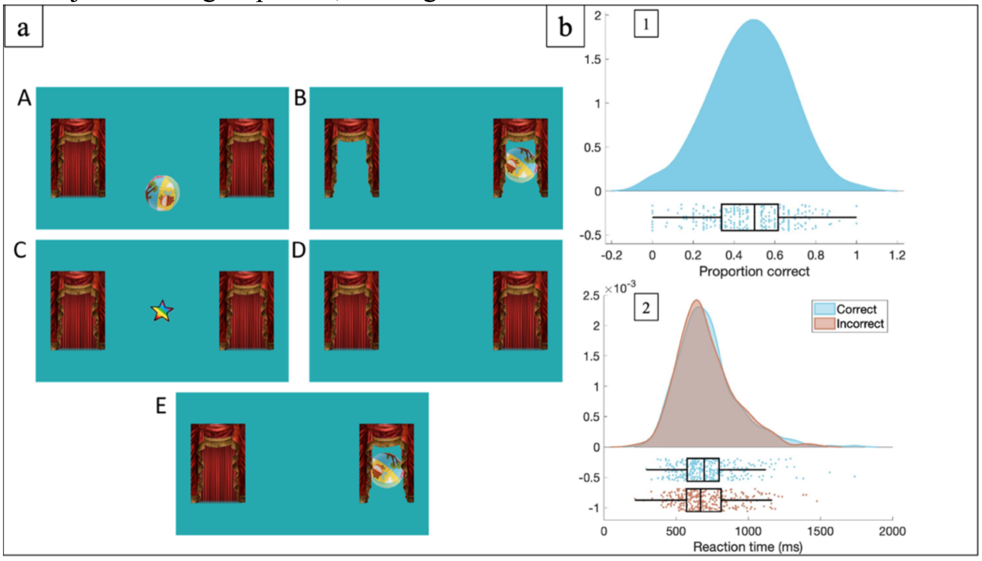


*Figure 5:* (a) Task figure and (b) raincloud plot of (b1) the proportion of correct trials and (b2) reaction times in correct and incorrect trials for the working memory task.

*2.2.5 Visual Search*

97.4% of children successfully completed the task with an average of 5 ($SD = 1$) valid trials per condition. Reaction times were slower in the conjunctive ($M = 1316.68$ms, $SD = 36.12$) than single feature displays ($M = 938.79$ms, $SD = 241.21$) with nine elements; (Overall effect of condition in ANOVA, $F(1.76, 450.90) = 159.90$, $p < 0.001$, $n = 257$, $\eta_p^2 = 0.38$; VS-S9-SRT versus VS-C9-SRT, $F(1, 256) = 255.46$, $p < 0.001$, $\eta_p^2 = 0.50$) and slower in the conjunctive thirteen ($M = 1391.81$ms, $SD = 372.28$) than conjunctive nine conditions; ($F(1, 256) = 5.62$, $p = 0.02$, $\eta_p^2 = 0.02$; Figure 6).

A similar repeated-measures ANOVA revealed there was also a difference in the proportion of correct trials across the three conditions; ($F(2, 680) = 309.11$, $n = 341$, $p < 0.001$, $\eta_p^2 = 0.48$). Pairwise comparisons revealed that the proportion of correct trials was higher in the singular nine condition ($M = 0.75$, $SD = 0.26$) versus the conjunctive nine conditions ($M = 0.45$, $SD = 0.28$); ($F(1, 340) = 374.47$, $p < 0.001$, $\eta_p^2 = 0.52$) and in the conjunctive nine versus the conjunctive thirteen conditions ($M = 0.38$, $SD = 0.26$); ($F(1, 340) = 19.17$, $p < 0.001$, $\eta_p^2 = 0.05$; Figure 6).

Results were consistent if children were only included with a minimum of 3 valid trials per condition (SM*2.3.2.5.*) whilst the same reaction time but not accuracy effects were found when accounting for trial variability, (SM*2.2.2.5.*). Thus, this task yielded good quantity and quality of data and found the expected condition effects when administered in a longer battery.

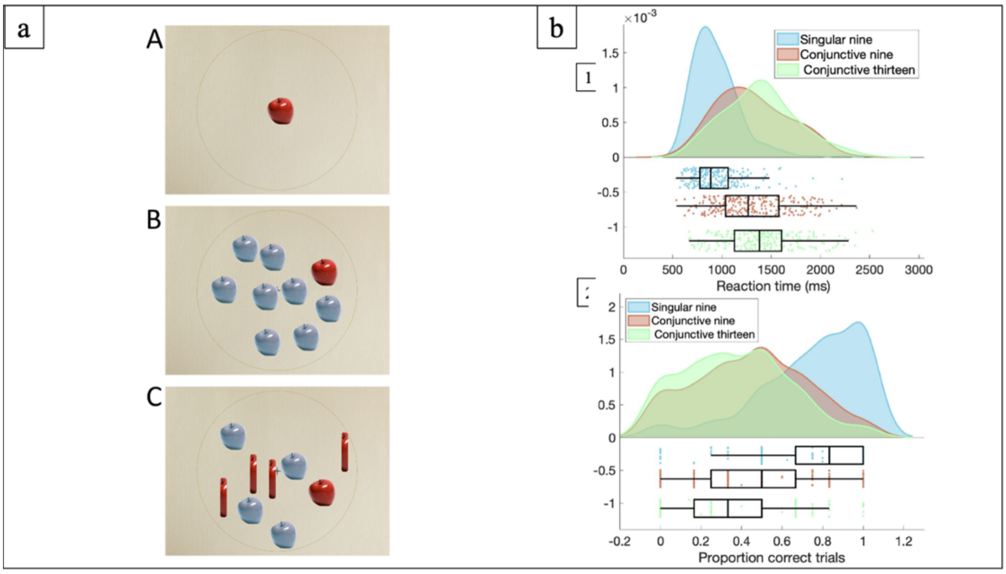


*Figure 6*: (a) Visual search task display and (b) raincloud plot of (b1) reaction times and (b2) accuracy to find the target in the singular nine, conjunctive nine and conjunctive thirteen conditions of the visual search task.

*2.2.6 Face pop-out*

73.1% of children successfully completed the task with an average of 7 (*SD* = 2) valid trials. Analysis showed more looking to faces than cars and noise for both duration of looking; (Overall, $F(1.39, 354.20) = 165.96$, $p < 0.001$, $\eta_p^2 = 0.39$; face versus car, $F(1, 255) = 60.31$, $p < 0.001$, $\eta_p^2 = 0.19$; face versus noise, $F(1, 255) = 607.29$, $p < 0.001$, $\eta_p^2 = 0.70$; means in table 3) and peak look, (Overall, $F(1.43, 355.21) = 94.38$, $p < 0.001$, $\eta_p^2 = 0.28$; face versus car, $F(1, 248) = 8.43$, $p < 0.001$, $\eta_p^2 = 0.03$; face versus noise, $F(1, 248) = 363.16$, $p < 0.001$, $\eta_p^2 = 0.59$; means in table S3, SM2.1.2).
A one-sample t-test revealed that the mean proportion of looking time to faces was significantly higher than chance, $t(255) = 14.01$, $p < 0.001$, CIs 95%[0.10, 0.14], $d = 0.86$, where chance level was 1 in 5 (0.2) and proportion of looking to faces was $M = 0.32$, $SD = 0.14$, figure 7. The proportion of trials in which the first look was to the face was also significantly higher than chance, $t(255) = 26.50$, $p < 0.001$, CIs[0.37, 0.43], $d = 1.67$, $M = 0.60$, $SD = 0.24$, figure 7. Results were consistent if children were only included with a minimum of 3 valid trials (SM*2.3.2.6.*), whilst peak look duration but not proportion looking was significantly impacted by data duration (SM*2.2.3.1.*). Thus, the pop-out task had a reasonable quantity and quality of data and as expected showed higher face orienting and face attention than other comparative stimuli.

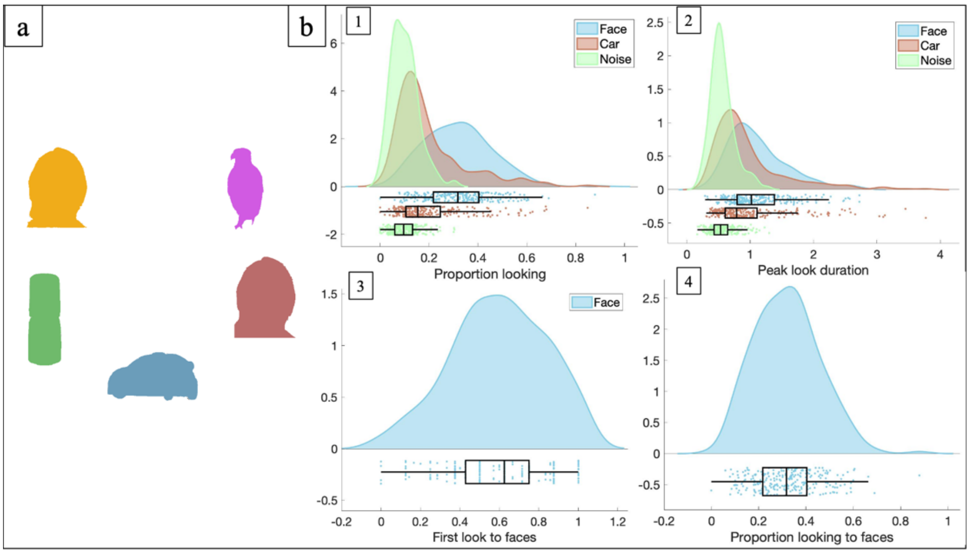


*Figure 7*: (a) Face pop-out task display and (b) raincloud plot of (b1) percentage looking and (b2) peak look duration to faces, car and noise, and (b3) proportion of trials in which the first look was to faces and (b4) percentage looking time to faces in the pop-out task.

2.1.2. Face Pop-out Task

Table S3: *Mean, standard deviation and n for overall looking, peak look and mean look duration to face, car and noise AOIs in the pop-out task*

| | | *Mean* | *SD* | *N* |
|---|---|---|---|---|
| Overall looking | Face | 0.32 | 0.14 | 256 |
| | Car | 0.20 | 0.14 | 256 |
| | Noise | 0.10 | 0.06 | 256 |
| Peak look duration | Face | 1.13 | 0.48 | 249 |
| | Car | 0.98 | 0.59 | 249 |
| | Noise | 0.56 | 0.19 | 249 |

*2.2.7 Dancing ladies*

71.1% of children successfully completed the task with an average of 4 (*SD* = 1) valid trials in each of the scrambled and social conditions.

An 2 x 2 repeated-measures ANOVA showed that peak look to faces ($M = 0.62$, $SE = 0.02$) was significantly higher than to objects ($M = 0.51$, $SE = 0.01$), ($F(1, 241) = 24.93$, $p < 0.001$, $\eta_p^2 = 0.09$), and peak look to faces was significantly greater in social ($M = 0.75$, $SD = 0.41$) than scrambled ($M = 0.49$, $SD = 0.27$), ($t(242) = 11.36$, $p < 0.001$, $d = 0.75$), figure 8. Proportion looking to faces ($M = 0.14$, se = 0.01) was also significantly higher than to objects ($M = 0.10$, se = 0.002), ($F(1, 248) = 29.66$, $p < 0.001$, $\eta_p^2 = 0.11$), and proportion looking to faces was significantly higher in social ($M = 0.20$, sd = 0.14) than scrambled conditions ($M = 0.07$, $SD = 0.06$), ($t(248)) = 16.67$, $p < 0.001$, $d = 1.21$). Other variables showed the same effect pattern (SM2.1.3.). Results were consistent if children were only included with a minimum of 3 valid trials of data (SM*2.3.2.7.*) whilst data duration significantly impacted some proportion looking and peak look duration measures, (SM*2.2.3.2.*).

Thus, this task yielded reasonable quantity and good quality of data and found the expected condition effects when administered in a longer battery.

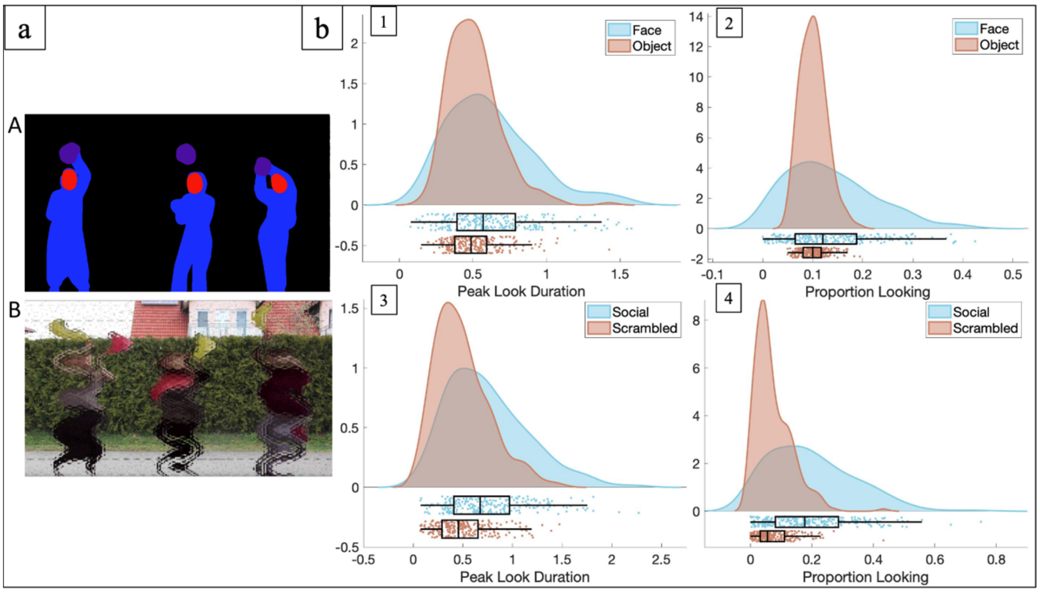


*Figure 8*: (a) Dancing ladies task display and (b) raincloud plot of (b1) peak look duration and (b2) proportion looking to faces and object across social and scrambled conditions, (b3) peak look duration and (b4) proportion looking time to faces in the social and scrambled conditions of the dancing ladies task.

In addition to key variables, other variables also attained from this task were number of looks to faces and mean look duration. Analyses with these variables indicated the same pattern as proportion looking and peak look duration, figure S1.

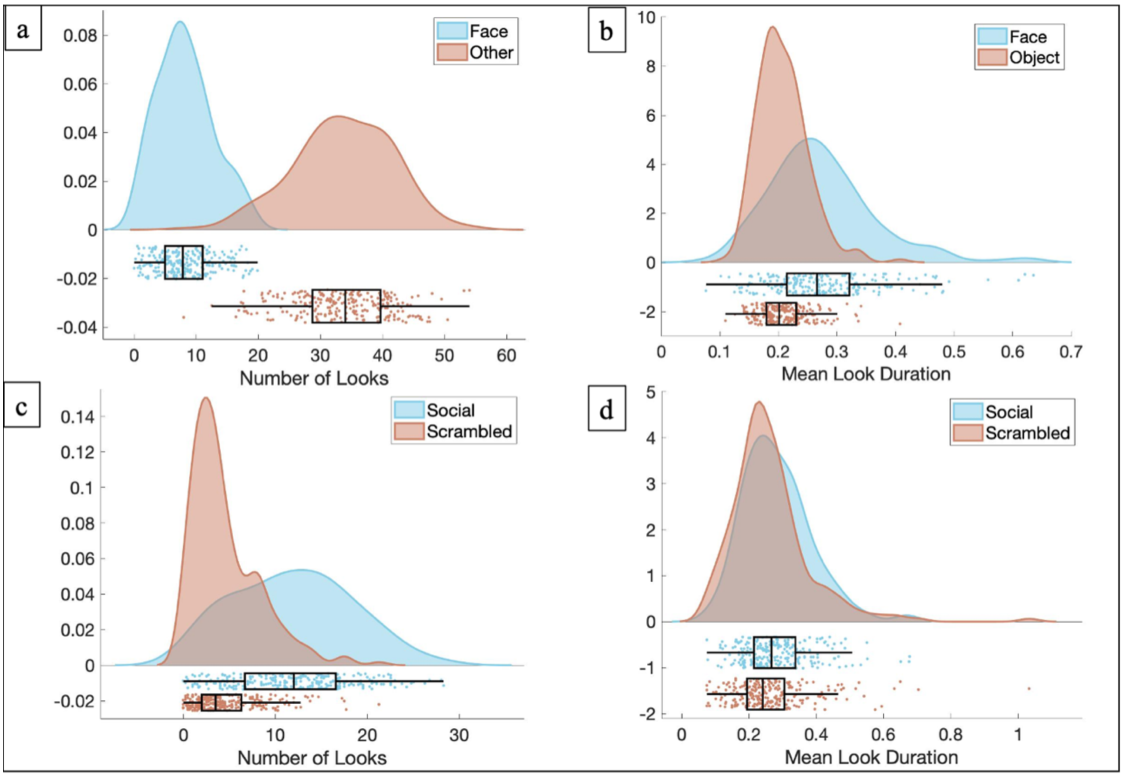


*Figure S1*: Raincloud plot of (a) number of looks and (b) mean look duration to faces and object across social and scrambled conditions, (c) number of looks and (d) mean look duration to faces in the social and scrambled conditions of the dancing ladies task

*2.2.8 Fifty faces*
70.3% of children successfully completed the task with an average of 71.9 (25) percentage valid data. A paired samples t-test showed that peak look to faces ($M$ = 3.65, $SD$ = 1.83) was significantly higher than to background people ($M$ = 0.46, $SD$ = 0.34), (t(226) = 25.95, $p$ < 0.001, $d$ = 2.42). Proportion looking to faces ($M$ = 0.57, $SD$ = 0.15) was also significantly higher than to background people ($M$ = 0.03, $SD$ = 0.03), ($t$(245) = 53.83, p < 0.001, $d$ = 4.99), figure 9. Results were consistent if children were only included if they provided at least 80% of valid data (SM*2.3.2.8.*) whilst data duration significantly impacted both measures (SM*2.2.3.3.*).
Thus, this task yielded reasonable quantity and quality of data and the expected condition effects when administered in a longer battery.

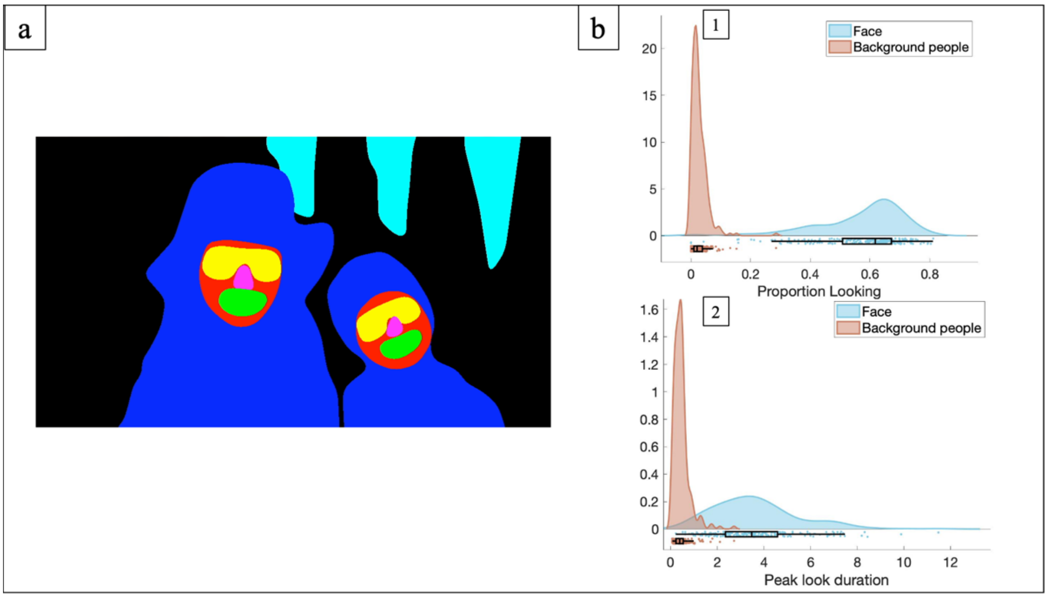


*Figure 9:* (a) Fifty faces task display and (b) raincloud plot of (b1) proportion of looking and (b2) peak look duration to faces and background people in the fifty faces task.

#### *2.2.2. Cut-off analyses*

For each task, participants were excluded according to cut-off criteria (Table S22); condition effect analyses as reported in the main text were then repeated. All statistical tests were conducted as in the main text; results are presented in tables only here for succinctness.

Table S22: *Exclusion criteria which were applied to each task prior to cut-off analyses being performed*

| Task | Cut-off criteria |
|---|---|
| Gap | < 5 valid trials per condition |
| Non-social Contingency | < 5 valid trials per condition |
| Reversal Learning | < 2 valid trials per phase |
| Working Memory | < 10 valid trials |
| Visual Search | < 3 valid trials per condition |

| | |
|---|---|
| Face pop-out | < 3 valid trials |
| Dancing Ladies | < 3 valid trials |
| Fifty Faces | < 20% trials valid |

*2.2.3. Trial level analyses in trial-based tasks*

Trial level analyses were conducted in trial-based tasks to assess whether condition effects were found when using data from each participants' individual trials (rather than an average of these per participant). We used the lmerTest package (Kuznetsova et al., 2017) in R (R Core Team, 2020) to conduct linear mixed models which enabled this variability to be included. We utilised a linear mixed-effects model rather than a traditional repeated-measures method as it allowed us to include data from multiple trials for each participant and thus to account for trial variance between subjects.

For each of the analyses outlined in table S6, a linear mixed effects analysis was performed with participant number entered as a random effect and condition as a fixed effect. A random intercept model was used; the variance-covariance structure was not specified.

Table S6*: Overview of linear mixed effect analyses*

| Task | Variable | Comparison |
|---|---|---|
| Gap-overlap | Reaction time | Baseline vs Gap, Baseline vs Overlap |
| Non-social contingency | Reaction time without zone-outs,<br>Reaction time to return to fixation stimulus | Sixty vs Zero, Sixty vs Hundred |
| Reversal learning | Reaction time | Learning vs reversal phase |
| Working Memory | Reaction time | Correct vs incorrect |
| Visual Search | Reaction time,<br>Proportion of correct trials | Conjunctive nine vs singleton nine,<br>conjunctive nine vs conjunctive thirteen |

*2.2.4. Data duration analyses in free viewing tasks*

For the following free-viewing tasks (face pop-out, dancing ladies & fifty faces), the duration of valid data from each task was controlled for as a measure of data quality. For the dancing ladies task separate data duration values were attained for each of the social and scrambled conditions. Data duration variables were centered around the mean and linear mixed analyses were conducted using the lmerTest package (Kuznetsova et al., 2017) in R (R Core Team, 2020). Participant number was entered as a random effect and data duration and AOI were fixed effects. For the dancing ladies task, separate data duration values were attained for each of the social and scrambled conditions and condition was also a fixed effect. A random intercept model was used; the variance-covariance structure was not specified. An ANOVA deviance test was conducted on each model; results from these are in table S1.

[[insert table here]]

Table S1: Results from condition comparisons for all tasks including (1) all participants who provided usable data, (2) participants after exclusion criteria (Table S?), (3) trial-level analyses and (4) data duration analyses
*
**
****This value is* $X^2$ *for the main effect of condition in an ANOVA deviance test conducted on the model*
